# Nanofitin®-Engineered Affinity Chromatography for Marker-Defined Extracellular Vesicle Enrichment in Scalable Downstream Processing

**DOI:** 10.64898/2026.04.17.719239

**Authors:** L.F. Koch, C. Golibrzuch, F. Cortopassi, K. Breitwieser, T. Best, E Wuestenhagen, M.J. Saul

**Author notes:** **Correspondence:** Meike J. Saul.

## Abstract

Extracellular vesicles (EVs) are lipid bilayer-enclosed particles that mediate intercellular communication through the transfer of bioactive molecules. Their growing relevance in translational applications demands downstream purification workflows that are selective, scalable, and compatible with robust impurity control. Conventional EV isolation methods primarily rely on physicochemical properties such as size, density, or charge and therefore co-enrich overlapping EV fractions together with non-vesicular impurities. Here, we establish a Nanofitin®-based affinity chromatography workflow for selective enrichment of a CD81-positive EV fraction under EV-compatible elution conditions. Nanofitin® candidate NF06 was identified by ribosome display against the large extracellular loop of CD81 and combined nanomolar affinity with favorable release behavior while retaining binding after repeated regeneration cycles. Static screening with recombinant CD81 and HEK293-derived EVs identified 1 M arginine at pH 10 as the most suitable elution condition. Dynamic chromatography on a 1 mL column using tangential flow filtration-concentrated HEK293 conditioned medium achieved 66.9% overall recovery with an elution step yield of 57.7%. In parallel, dsDNA, host cell protein, and total protein were reduced by 2 to 3 log relative to conditioned medium. Nano flow cytometry showed enrichment of the CD81-positive EV fraction from 40% in conditioned medium to more than 90% in the eluates, together with a smaller and narrower particle size distribution. These results demonstrate that Nanofitin®-based affinity chromatography provides a practical route toward marker-defined EV enrichment that combines selective capture, EV-compatible release, and substantial impurity clearance in a chromatography-compatible process format.

## Introduction

Extracellular vesicles (EVs) are nano-sized particles enclosed by a lipid bilayer, secreted by all cell types into biological fluids. They transport various bioactive molecules and play crucial roles in intercellular communication, influencing both physiological and pathophysiological processes (1, 2). Beyond their biological relevance, EVs are increasingly explored for translational applications (e.g. drug delivery), which places stringent demands on scalable, reproducible, and well-controlled downstream processing and robust quality control. The term “EV” encompasses a heterogeneous population of subtypes that vary in biogenesis, composition, size, and function. Although EVs are commonly classified according to their biogenesis, overlapping formation mechanisms and shared physicochemical properties complicate clean separation using conventional workflows (1). This operational overlap motivates approaches that enable selective enrichment of marker-defined EV fractions rather than broad EV mixtures. The protein composition of EVs is cell-type specific and influenced by the physiological state of the originating cells, in addition to shared biogenic pathways (3). Tetraspanins (TPs), a family of proteins characterized by four transmembrane domains along with extracellular and cytoplasmic domains, are closely associated with EV formation (4, 5). In addition to their roles in endosomal networking and EV biogenesis, TPs have been implicated in cargo organization and sorting, supporting their widespread use as practical EV surface markers (4, 6, 7). Due to their pivotal role, TPs like CD9, CD63, and CD81 are commonly used as EV markers (8). CD81 is notably present on a wide range of EVs derived from various parental cells (9). CD81 also provides well-characterized extracellular domains that are accessible for affinity targeting and systematic binder selection. CD81 was chosen as an exemplary benchmark target to evaluate the complete development cycle from binder selection, matrix coupling, chromatographic capture/elution, and orthogonal analytical validation.

For translational and pharmaceutical applications, EV downstream purification must not only enrich vesicles but also deliver robust impurity clearance, process consistency, and scalability. Current isolation and concentration strategies, spanning centrifugation/filtration, density-based approaches, ultrafiltration, size exclusion chromatography (SEC), anion exchange chromatography (AEX), and tangential flow filtration (TFF) face trade-offs in yield, purity, productivity, and selectivity and frequently co-enrich non-vesicular impurities (10–14). Overall, approaches relying primarily on size, density, or charge inherently recover overlapping EV subtypes and do not directly enable surface-marker-defined enrichment. This creates a mismatch between translational requirements (marker-defined identity, reproducibility, impurity control) and separation principles dominated by bulk physicochemical properties, motivating affinity-based unit operations for marker-defined enrichment. An ideal downstream unit operation should combine (i) marker-selective capture, (ii) controllable and EV-compatible release, (iii) chromatography compatibility to support scalability and reproducibility, and (iv) quantifiable impurity clearance. To minimize variability among isolated subtypes, affinity-based purification methods can be employed to selectively enrich EVs expressing specific surface markers, achieving higher purity through targeted interactions. Conventional affinity chromatography typically utilizes antibody ligands for surface-marker-specific isolation; however, this approach is associated with increased production costs, limited lifespan, and harsh elution conditions that may denature or damage EVs (15, 16). These limitations motivate affinity ligands and chromatographic formats that retain marker selectivity while enabling robust column operation and EV-compatible, controllable release. Alternatives to antibodies, including RNA/DNA-based tools and smaller proteins or polypeptides, aim to address these challenges (17). In this context, small recombinant affinity ligands are particularly attractive for chromatography formats because they can support robust column operation, regeneration, and controlled release while maintaining marker selectivity.

Nanofitins®, derived from the ~7 kilodalton (kDa) DNA-binding protein Sac7d of the archaeal microorganism *Sulfolobus acidocaldarius*, provide a promising alternative (18). By generating a mutant Sac7d library and selecting affinity binders, a diverse range of targets can be addressed (19). As compact (~7 kDa), single-domain binders that are readily produced recombinantly, Nanofitins® enable reproducible ligand manufacture and straightforward engineering for controlled immobilization on chromatographic matrices. Their robust scaffold is well suited to stable column operation and regeneration. Moreover, selection campaigns can be directed toward binders that combine high-affinity capture with EV-compatible, controllable release, aligning ligand properties with chromatographic process requirements. By immobilizing engineered Nanofitin® ligands on a chromatography matrix, we create a programmable affinity surface whose selectivity is defined by the binder rather than bulk physicochemical properties. Nanofitins® have been successfully developed for several clinically relevant targets, serving as drug delivery agents or direct inhibitors (20, 21). Furthermore, they have been effectively utilized for the one-step purification of a Streptococcus vaccine (22). In principle, this also enables a modular “ligand-exchange” concept in which the same core process can be retargeted to distinct EV surface markers by exchanging the binder.

Here, we establish a Nanofitin®-based affinity chromatography workflow for selective enrichment of a CD81-positive EV fraction under EV-compatible elution conditions and benchmark performance metrics relevant to scalable downstream processing. Binders were selected against the large extracellular loop (LEL) of CD81 and screened for controlled target release under EV-compatible elution conditions. Selected Nanofitin® was coupled to Eshmuno® base beads, and binding/elution were optimized and validated using recombinant CD81 LEL and HEK293-derived EVs. The resulting EV-enriched fractions were characterized with orthogonal analytics to assess marker enrichment, size distribution, recovery, elution step yield, and chromatographic impurity clearance.

## Results

### Selection of a CD81-Binding Nanofitin® with Controlled Dissociation Properties

To generate binders against CD81, Nanofitin® selection (Fig. 1A) was performed using the recombinant large extracellular loop (LEL) of CD81, spanning Phe113 to Lys201 and fused to an Fc tag, to focus selection on the isolated CD81 extracellular domain rather than other EV surface components (Fig. 1C). Selection was carried out by ribosome display over four rounds with increasing stringency (Fig. 1A, B).

**Figure 1.**
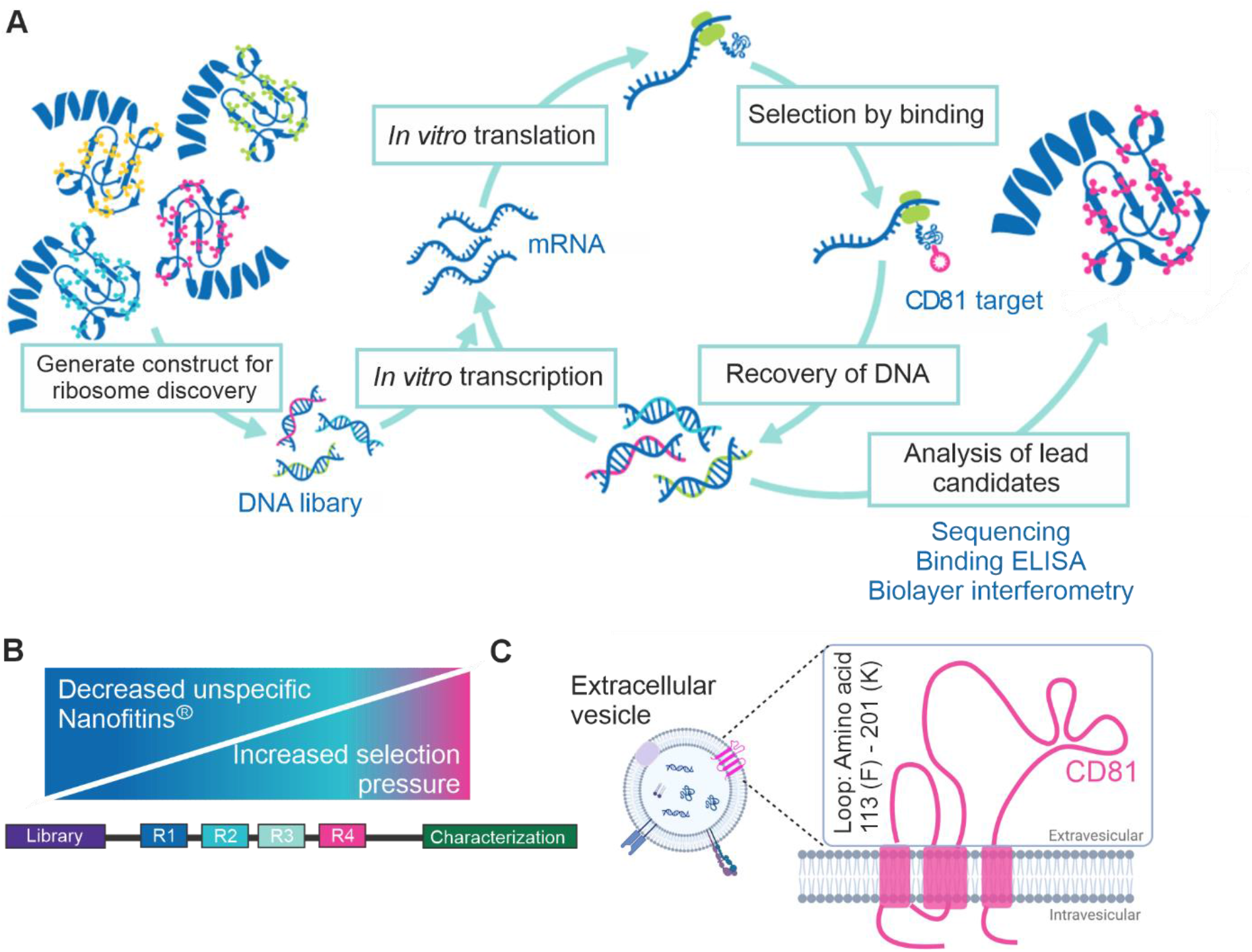
Selection process of Nanofitins® targeting CD81. (A) Illustrative representation of ribosome display cycle, starting with transcription of generated DNA Nanofitin® library and translation to form RNA-ribosome-Nanofitin® complex. Selection of Nanofitins® through binding of the complex with immobilized protein of interest. After dissociation, RNA of bound complexes is reverse transcribed, generating starting material for a new round of selection. After final selection rounds, lead candidates are characterized. (B) Schematic of Nanofitin® specificity throughout selection routes. Diversity of Nanofitin® binders reduces throughout selection rounds while selection pressure increases, resulting in the enrichment of CD81-binding Nanofitins®. (C) Presentation of target of interest, present on the surface of extracellular vesicles. CD81, a tetraspanin and transmembrane protein that contains two extracellular loops. Recombinant large extracellular loop (LEL), from Phe113 to Lys201, was used as proxy for the selection of Nanofitin® binders against CD81 due to off targets present on extracellular vesicle surface. (Figure created using BioRender.com)

Enrichment of CD81-binding Nanofitins® from the naïve library was monitored by Enzyme-linked Immunosorbent Assay (ELISA) using pooled binders from each selection round. An increase in CD81-specific relative to unspecific binding was observed from round two onwards, indicating progressive enrichment of CD81-binding candidates (Sup.Fig.1). After completion of the selection, 95 GFP-fused clones were picked from fluorescent *Escherichia coli* expression cultures representing the final selection round. Initial characterization was performed by CD81-targeting ELISA from cell lysate and showed a broad range of CD81 binding signals across the clone set (Fig. 2A). Based on this primary screening, 11 candidates were purified and further characterized by ELISA (Fig. 2B) as well as biolayer interferometry (BLI).

**Figure 2.**
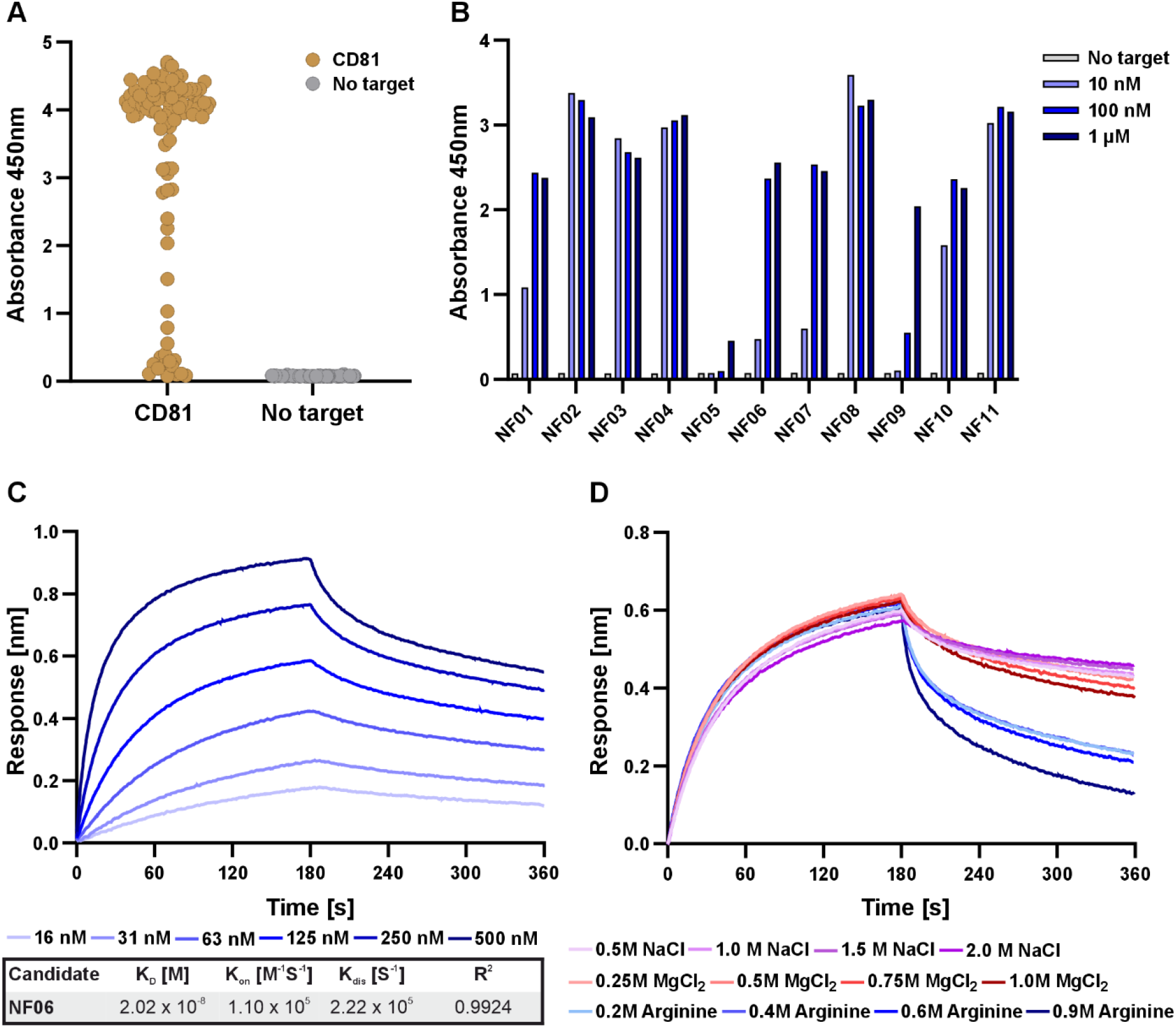
Selection and characterization of CD81-binding Nanofitin® candidates from ribosome display. (A) CD81 binding screening in crude lysate of 95 picked candidates from library after cloning of last ribosome display selection round, quantified by enzyme linked immunosorbent assay (ELISA) and compared to unspecific binding. (B) Screening of binding characteristics of 11 purified candidates in three different concentrations against CD81 and no target, quantified by ELISA. (C) Dose-dependent binding response (layer thickness) and calculated affinity parameters using 1:1 stoichiometry fit model of best performing candidate NF06, measured using Biolayer interferometry (BLI). (D) Screening of dissociation conditions of NF06 from CD81, including the addition of NaCl, MgCl_2_ and arginine in 50mM TRIS buffer at pH 10 (NaCl, arginine) or pH 9 (MgCl_2_), measured using BLI. Data represents single measurement.

Binding dynamics of the purified candidates were evaluated by concentration titration from 10 nM to 1 µM. NF06 displayed a dose-dependent increase in binding with a strong overall signal, whereas NF02 and NF08 showed high but largely concentration-independent signals and NF05 showed only weak binding. The five most promising variants were produced with terminal cysteine instead of GFP to enable site-specific conjugation. After biotinylation of the ligand, BLI measurements were performed to assess CD81 binding and release under the tested elution conditions. Among these five candidates, NF06 showed the most favorable combination of binding behavior, expression level, and elution profile and was therefore advanced as the lead ligand for subsequent characterization.

BLI titration of CD81 from 16 to 500 nM yielded a dissociation constant (K_D_) of 2.02 × 10^-8^ M for NF06 (Fig. 2C). Elution capability of NF06 was screened under alkaline conditions with the addition of either NaCl, MgCl_2_, or arginine (Fig. 2D). Whereas NaCl and MgCl_2_ did not measurably affect dissociation across the tested concentration range, arginine induced a concentration-dependent increase in dissociation. After identification of NF06 as a suitable CD81-binding candidate with release under the tested elution conditions, its stability towards caustic and low-pH conditions was assessed by BLI (Sup.Fig.2). After 20 cycles in 0.1 M NaOH or 0.1 M glycine pH 2.0, NF06 showed only minimal loss of CD81 binding.

### Immobilized NF06 Retains Binding and Defines an EV-Compatible Elution Window

Following lead selection, NF06 was immobilized on Eshmuno® base beads to evaluate its performance in a chromatographic format. Static binding capacity, SBC, assays were first performed with recombinant CD81 LEL to confirm the coupling procedure did not impair ligand function. Flow-through quantification showed reduced CD81 concentration after contact with the resin, corresponding to an SBC of 3.55 mg/mL resin (Sup.Fig.3, A). This confirmed that immobilized NF06 retained its binding capability towards recombinant CD81. The elution behavior of resin-coupled NF06 was then assessed using recombinant CD81 LEL to identify suitable arginine concentration, pH, and possible effects of NaCl addition. Increasing the arginine concentration from 0.1 M to 1 M resulted in increased elution, reaching 1.50 mg/mL resin (Sup.Fig.3, B) Supplementation of 1 M NaCl to 1 M arginine further increased elution to 1.65 mg/mL resin. In contrast, acidic elution buffers such as 1 M arginine at pH 4 or 0.1 M glycine at pH 3 did not outperform alkaline elution conditions. Elution with wash buffer alone did not release detectable amounts of CD81, confirming retention on the resin.

To confirm that NF06 also binds native CD81 on the EV surface, static binding assays were repeated using purified HEK293-derived EVs. Flow-through (FT) quantification by CD81 ELISA gave an SBC of 1.94 × 10^11^ particles/mL resin (Sup.Fig.3, C). This corresponded to 32% of total loaded EVs and 96% of loaded CD81-positive EVs. Elution screening was then repeated with EVs and revealed a different pattern from that observed for recombinant CD81 LEL (Sup.Fig.3, D). As observed for recombinant CD81, increasing arginine concentration up to 1 M at pH 10 improved elution and resulted in the highest EV recovery. However, addition of 1 M NaCl reduced elution to 59% relative to 1 M arginine at pH 10 alone. Acidic elution at pH 4 also reduced recovery to 63% relative to 1 M arginine at pH 10. Control conditions with 1 M arginine at pH 7.4 or pH 10 without arginine gave lower relative recoveries of 59% and 40%, respectively.

Additional elution conditions were tested to further improve EV recovery (Sup.Fig.3, E). Neither 1.5 M arginine, 0.1% Tween-20, nor 1 M MgCl_2_ improved elution compared with 1 M arginine at pH 10. Increasing arginine concentration to 1.5 M at pH 4 improved recovery compared with 1 M arginine at pH 4 but remained below the performance of 1 M arginine at pH 10. Addition of MgCl_2_ under acidic conditions further reduced elution. Based on these results, 1 M arginine at pH 10 was selected for the subsequent column experiments.

### Dynamic NF06 Affinity Chromatography Enriches CD81-Positive EVs While Reducing Process Impurities

The NF06-based affinity resin and the selected elution conditions were next evaluated in a complete chromatography workflow using TFF-concentrated HEK293-conditioned medium as load. Duplicate runs were performed on independent 1 mL prototype columns to assess process performance and reproducibility. Chromatographic steps were monitored inline by UV detection at 280 nm and multi-angle light scattering, MALS, and the corresponding fractions were collected for further analysis (Fig. 3, Fig. 4A).

**Figure 3.**
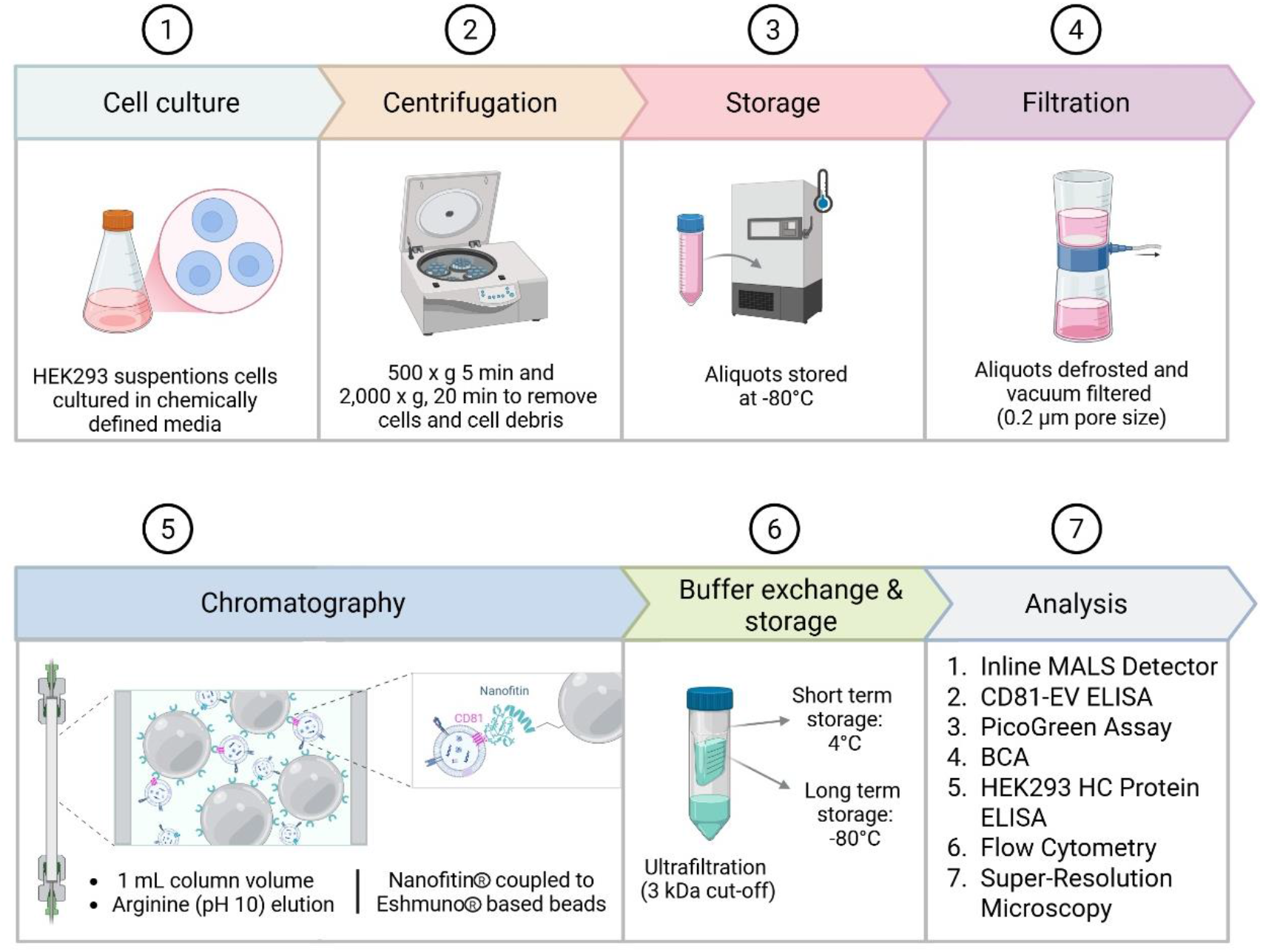
Workflow of Nanofitin®-based affinity chromatography for selective enrichment of CD81-positive extracellular vesicles (EVs) from cell culture supernatant. The process begins with EV production from suspension-adapted HEK293 cells cultivated in chemically defined medium. Cells and cellular debris are removed by centrifugation, and conditioned medium (CM) containing EVs is aliquoted and stored at −80 °C. After thawing, CM is concentrated 10-fold by tangential flow filtration (TFF) and loaded onto a 1 mL anti-CD81 Nanofitin® Eshmuno® prototype column. EVs are eluted with 1 M arginine at pH 10 and, after buffer exchange, characterized using orthogonal analytical methods. (Figure created using BioRender.com)

**Figure 4.**
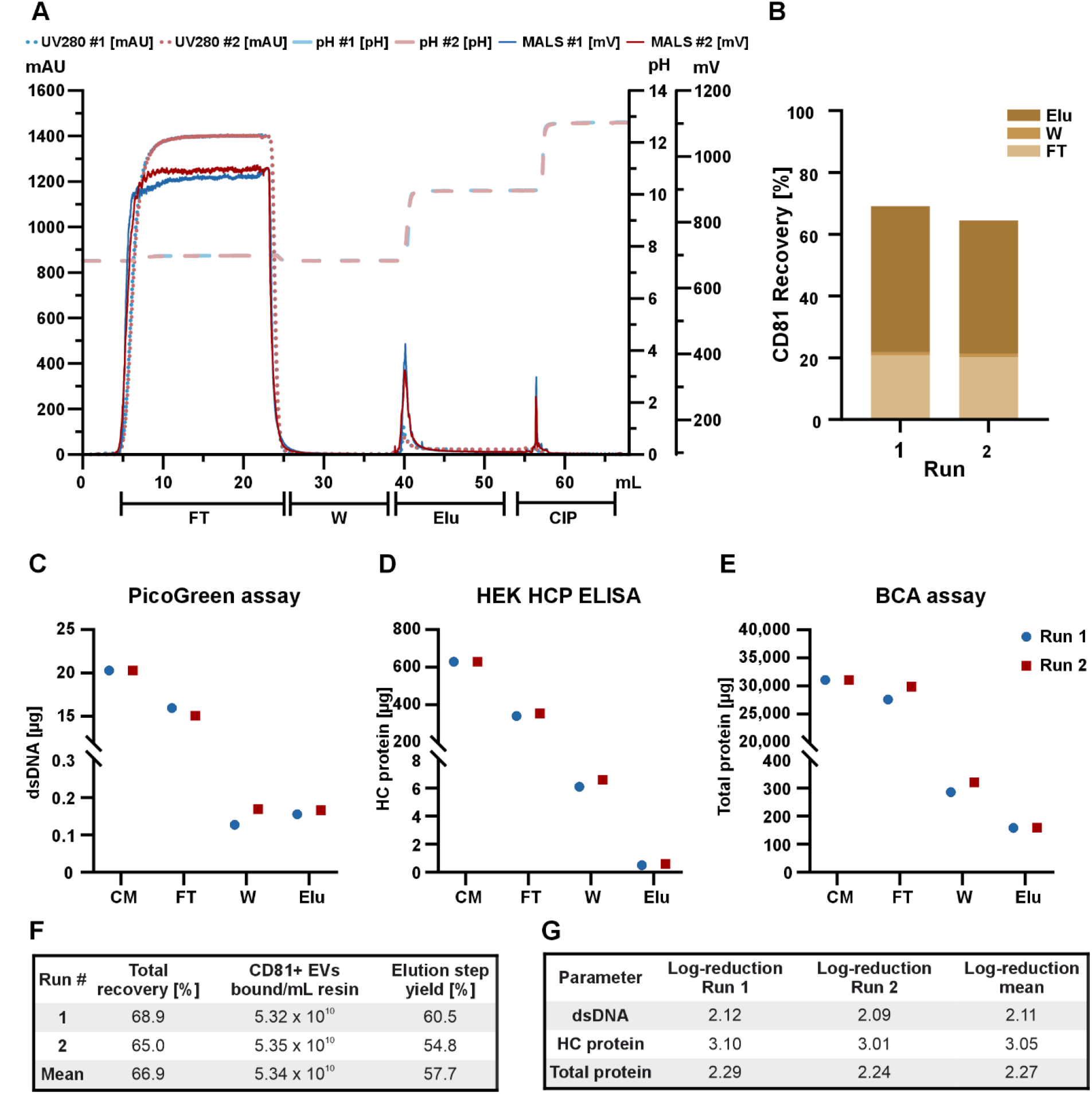
Performance characterization of CD81-targeted affinity chromatography for extracellular vesicle (EV) purification and impurity clearance using Nanofitin® NF06 prototype column. (A) Chromatogram overlay of two separate runs with individual prototype columns, run 1 (blue), run 2 (red). Depict are chromatographic steps, Flow-through (FT), Wash(W), Elution (Elu), Cleaning-In-Place (CIP) which are monitored using UV detector at 280nm (dotted line), Multi-angle light scattering (MALS) detector (continuous line) and pH probe (dashed line). (B) Cumulative CD81-positive EV recovery for the chromatographic steps FT, W and Elu based on loaded EV amount (100%). (C) Quantification of double stranded DNA (dsDNA) using PicoGreen assay present in conditioned medium (CM) as Load, FT, W and Elu. (D) Quantification of host cell proteins (HCP) using HEK HCP ELISA present in CM as Load, FT, W and Elu. (E) Quantification of total protein using bicinchoninic acid (BCA) assay present in CM as Load, FT, W and Elu. (F) Table showing total recovery, CD81-postive EV binding capacity and elution step yield for both individual runs. (G) Table showing log reduction of impurities dsDNA, HC protein and total protein for both individual runs. Data represents mean of two technical replicates.

Real-time MALS monitoring revealed a continuous signal during the loading phase in both runs (Fig. 4A). During elution with 1 M arginine at pH 10, the MALS signal increased again, indicating release of retained particles. A final MALS peak was observed during cleaning-in-place, CIP. The UV280 signal followed the same general trend during loading and elution, while no relevant signal was detected during CIP. Quantification of CD81-positive EVs across all chromatographic fractions showed a mean overall recovery of 66.9% (Fig. 4B, F). The largest recovered fraction was found in the elution pool, corresponding to 45.3% of the loaded material, while 20.5% and 1.1% were recovered in the flow-through, FT, and wash fractions, respectively. This resulted in a mean binding of 5.34 × 10^10^ CD81-positive EVs per mL resin. Based on the bound particles, the mean elution step yield from both runs was 57.7%, with a particle concentration of 8.08 × 10^10^ p/mL in the pooled eluate.

Impurity clearance was evaluated for double-stranded DNA (dsDNA), host cell proteins (HCP) and total protein (Fig. 4C-E, G). All three impurity classes decreased across the chromatographic steps and reached their lowest levels in the elution fraction. Compared to conditioned medium, the eluate showed a 2.11-log reduction in dsDNA, a 3.05-log reduction in HCP, and a 2.27-log reduction in total protein. Additional Asymmetric-flow field-flow fractionation (AF4) analysis further assessed protein and particle distributions in conditioned medium and eluate (Sup.Fig.4). In conditioned medium, an early UV280 peak was detected separately from the later particle peak. In the elution sample, no separate early protein peak was observed and the remaining UV280 signal co-eluted with the particle fraction.

The presence of CD81-positive EVs in the pooled FT prompted analysis of subsequent FT fractions (Sup.Fig.5). After an initial increase in the first FT fraction, the CD81-positive EV signal remained at approximately 23% of the load until completion of the loading phase. A progressive increase toward feed concentration was not observed under the tested conditions.

### Affinity-Enriched Eluates Display a Narrowed, CD81-Enriched EV Population and Remain Stable under Elution-Relevant Conditions

To assess the impact of CD81-targeted affinity chromatography on EV populations present in conditioned medium, the pooled elution fractions were characterized and compared with the feed. Particle size distribution was first analyzed by nano flow cytometry (Fig. 5A-C). From the resulting size distributions, mean diameter, D10, D50, D90, and Span were calculated (Fig. 5D). While D10 differed only marginally between conditioned medium and eluate, with values of 66.0 nm and 65.3 nm, larger differences were observed for the remaining particle parameters. D50 decreased from 81.0 nm in conditioned medium to 72.3 nm in the eluate, and D90 decreased from 110.5 nm to 90.5 nm. Consistent with these shifts, the Span decreased from 0.56 in conditioned medium to 0.34 and 0.36 in the two eluate samples. The mean particle diameter was also lower in the eluates than in conditioned medium, with values of 76.0 nm and 76.7 nm compared with 85.7 nm.

**Figure 5.**
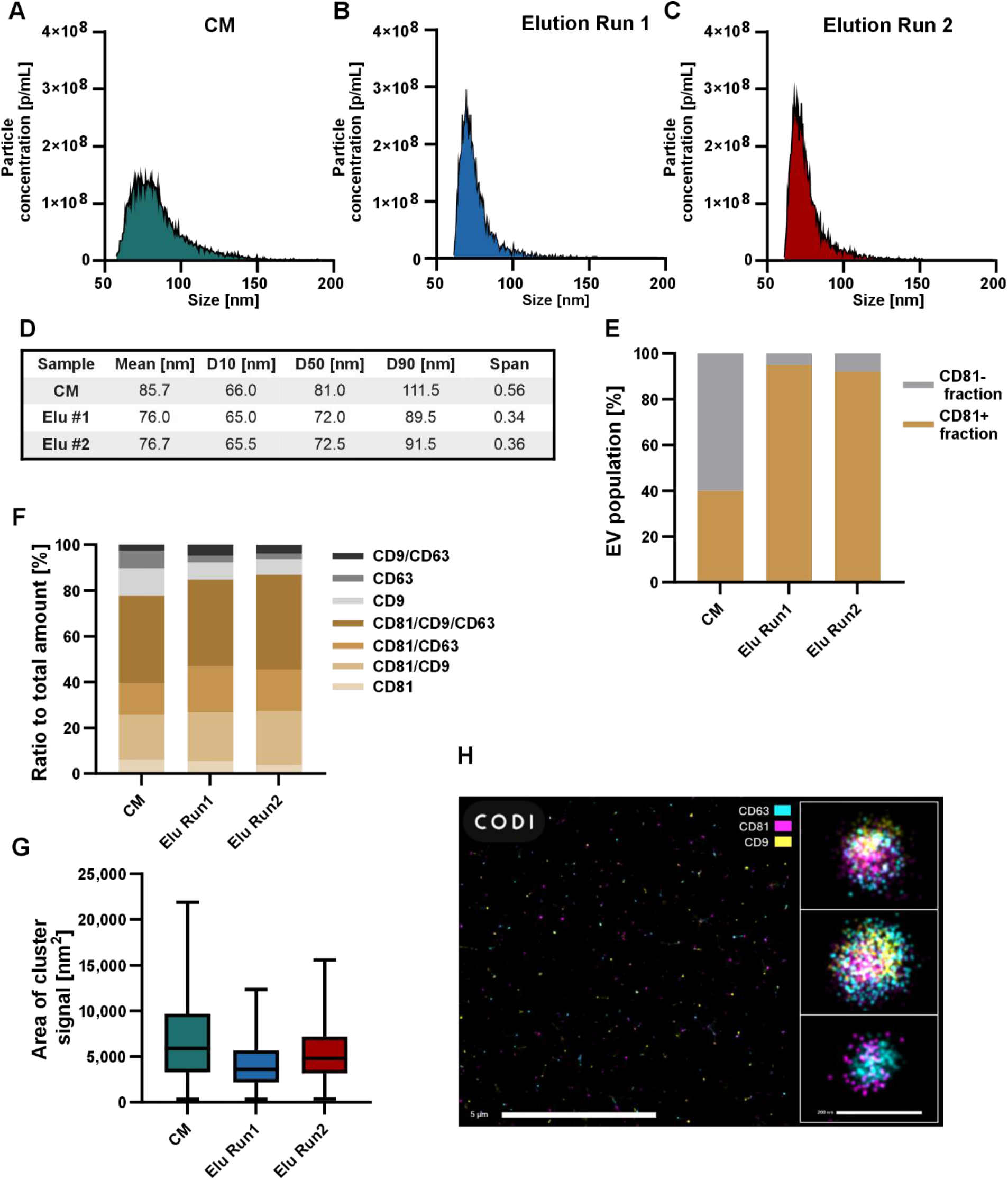
Characterization of affinity-enriched extracellular vesicles by size distribution, CD81 enrichment, and tetraspanin composition. (A-C) Particle size distribution of conditioned medium (CM) and eluate fractions from NF06 affinity chromatography runs 1 and 2, measured by nano flow cytometry. (D) Calculated size distribution parameters mean, D10, D50, D90, and Span for CM and eluate fractions from runs 1 and 2. (E) Fraction of CD81-positive particles in CM and eluate fractions from runs 1 and 2, measured by immunofluorescence staining and nano flow cytometry. Data represent the mean of two technical replicates. (F) Colocalization of the tetraspanins CD9, CD63, and CD81 in EVs from CM and eluate fractions from runs 1 and 2, measured by fluorescence staining and super-resolution microscopy. (G) Fluorescence cluster area derived from the super-resolution microscopy data shown in panel F with outlier removal (Q=1). Box represents 25^th^ to 75^th^ percentile, line represents median, and whiskers cover min to max. Data represent the mean ± standard error of the mean (SEM) of 5 technical replicates. (H) Exemplary Super-Resolution Microscopy (dSTORM) overview and zoom-in pictures of immunofluorescent stained EVs from the Elution of NF06 affinity chromatographic run 1, CD9 (yellow, 488 nm), CD63 (blue, 561 nm), CD81 (magenta, 647 nm).

The observed size shift was further assessed by AF4 analysis (Sup.Fig.4). In conditioned medium, the particle peak eluted later than in both eluate samples. This difference was consistent with the smaller particle size distribution observed in the affinity-enriched fractions. To evaluate enrichment of the target-positive population, samples were stained for CD81 and analyzed by Nanoflow Cytometry (Fig. 5E). In conditioned medium, 40% of detected particles were CD81-positive, whereas 60% were CD81-negative. After affinity chromatography, the CD81-positive fraction increased to 95% in elution sample 1 and 91% in elution sample 2, while the corresponding CD81-negative fractions decreased to 5% and 9%, respectively. These data confirmed selective enrichment of a CD81-positive EV fraction by Nanofitin® affinity chromatography. To assess the membranous nature of particles present in conditioned medium and eluate, samples were treated with Triton™ X-100 and quantified by nano flow cytometry (Sup.Fig.6). After Triton™ X-100 treatment, particle counts in conditioned medium were reduced by 83.6%, while the eluate fractions from run 1 and run 2 were reduced by 90.6% and 87.9%, respectively. Next, colocalization of the TPs CD9, CD63, and CD81 was analyzed in EVs from conditioned medium and both eluate fractions using super-resolution microscopy (Fig. 5F, G; Sup.Fig.7). In CM, CD81 was the most abundant TP, with a cumulative ratio of 77.7% between single (6.0%), double (33.5%) and triple (38.2%) positivity. CD9 (12.0%) and CD63 (7.7%) and their colocalization (2.7%) were present to a much lesser extent. While CD81 was already the most abundant TP in CM, Elution samples from run 1 and 2 showed a further increase in cumulative CD81 ratio to 84.8% and 86.9%, respectively. Cumulative increase of CD81 ratio was driven by increase in CD81/CD9 (21.4 - 23.8%) and CD81/CD63 (20.3 - 18.0%) double-positive EVs, whereas triple-positive ratio remained similar (37.8 - 41.4%). Increase in CD81 ratio was accompanied by reduction of both CD9 (7.4 - 6.9%) and CD63 (3.0 – 2.3%) single positive EVs Calculation of the area of fluorescent clusters (Fig.5G) revealed a decrease in both range of measured cluster area, as well as median cluster area (5906 nm^2^ CM,3614 nm^2^ Elu Run1, 4824 nm^2^ Elu Run2).

To evaluate EV stability under elution-related conditions, purified EVs were incubated in buffers containing 0.1 M to 1 M arginine at pH 10 for up to 24 h at 4 °C and were then quantified by nano flow cytometry (Sup.Fig.8). After normalization to the corresponding PBS control at each time point, no decrease in particle concentration was observed for any of the tested arginine concentrations in the current data set.

## Discussion

This study establishes a Nanofitin®-based affinity chromatography workflow for selective enrichment of a CD81-positive EV fraction under EV-compatible elution conditions. The central technical advance is not only the identification of a CD81-binding ligand, but the integration of binder selection, release screening, chromatography-compatible immobilization, and quantitative process evaluation into one coherent purification strategy. In this framework, NF06 combined nanomolar target binding with favorable release behavior in the presence of arginine and retained binding after repeated exposure to both caustic and low-pH regeneration conditions. Under dynamic operation, this translated into selective enrichment of CD81-positive EVs together with substantial reduction of dsDNA, HCP, and total protein. These findings support Nanofitin®-based affinity capture as a technically relevant unit operation for marker-defined EV enrichment in translation-oriented downstream processing.

CD81 was chosen as a practical benchmark because it is readily detectable on HEK293-derived EVs, accessible for binder selection through its extracellular loop, and sufficiently abundant for quantitative chromatographic evaluation(9, 23). At the same time, CD81-based enrichment should not be equated with purification of a biologically homogeneous EV subtype. MISEV2018 and its recent update emphasize that EV is an operational term and that marker-based enrichment alone does not justify assignment to a defined biogenesis class without additional mechanistic evidence (24, 25). This caution is consistent with work showing that TP-positive EVs are not biologically uniform and that CD81 does not map onto a single vesicle category or trafficking route (26, 9). Accordingly, the present workflow should be interpreted as selective enrichment of a CD81-positive EV fraction rather than purification of one discrete EV subtype.

A key finding of this study is that binder selection and elution design have to be developed together when the target is an intact vesicle rather than a soluble protein. Initial selection against recombinant CD81 LEL was necessary to avoid enrichment of binders against unrelated EV surface components. However, the subsequent comparison between recombinant CD81 and native EVs showed that favorable elution behavior on the isolated target did not translate directly to the vesicular context. NaCl modestly improved elution of recombinant CD81 but reduced EV recovery, and acidic conditions were likewise unfavorable in the EV setting. These differences show that EV purification cannot be optimized solely on the basis of soluble target behavior. Elution therefore has to be evaluated directly on intact vesicles. This is consistent with literature showing that pH and buffer composition can affect apparent EV stability and colloidal behavior, including shifts in particle concentration and zeta potential under acidic or altered ionic conditions (27, 28). The identification of 1 M arginine at pH 10 as the best-performing elution condition is therefore one of the central technical outcomes of the study. Arginine is well established in affinity chromatography as an eluent that can facilitate dissociation of affinity complexes and, in protein systems, can reduce aggregation relative to more conventional acidic strategies (29– 31). Although EV capture is not directly analogous to antibody-Protein A chromatography, the same process principle applies. Elution should weaken the interaction while minimizing collateral effects on the affinity-enriched EV fraction (32). In the present work, alkaline arginine elution emerged as the best compromise between release efficiency and EV compatibility. The current arginine stability experiment supports this conclusion at the level of particle counts, but stronger claims regarding preservation of vesicle integrity or biological activity should remain linked to the Triton™-based membrane disruption assay and the additional orthogonal characterization data.

The dynamic chromatography data highlight both the current strengths of the workflow and the main remaining optimization levers. The overall recovery of 66.9% and elution step yield of 57.7% provide a solid starting point for a first-generation CD81-targeted affinity process. At the same time, the difference between static EV binding and dynamic column performance indicates that not all CD81-positive EVs were captured efficiently under flow. In the static format, 96% of CD81-positive EVs were retained, whereas under dynamic conditions 20.5% remained in the FT. Subsequent FT analysis did not show the progressive increase expected for classical breakthrough but instead plateaued at approximately 23% of the feed concentration. These observations are consistent with incomplete capture of a CD81-positive subpopulation under the applied flow conditions rather than simple saturation of the resin. Possible contributors include residence time, mass transfer limitations, ligand accessibility on the resin, and heterogeneity in CD81 presentation on EV surfaces.

In parallel, incomplete desorption is indicated by the difference between bound and eluted material and by the residual signal detected during CIP. Unspecific EV losses during handling and chromatographic processing have been described previously (33, 14), but the present data suggest that incomplete recovery cannot be attributed to a single mechanism. Instead, both capture efficiency and release efficiency are likely to contribute under the current conditions. Together, these findings define a clear process-development space involving residence time studies, ligand density optimization, and refinement of the elution profile.

Beyond recovery, one of the strongest features of the workflow is the associated impurity clearance. Relative to conditioned medium, the eluate showed reductions of 2.11 log for dsDNA, 3.05 log for HCP, and 2.27 log for total protein. This is an important aspect of translation-oriented EV downstream processing, because selective enrichment alone is not sufficient if process-related impurities remain poorly controlled. In the present study, impurity reduction was achieved together with marker-defined enrichment in a single affinity step, which strengthens the practical relevance of the workflow beyond capture selectivity alone.

Affinity-based EV purification remains a relatively small but growing field, and cross-study comparison is inherently limited by differences in feedstock composition, target definition, analytical readouts, and process format. Peptide-based affinity chromatography has emphasized the importance of balancing selective capture with elution conditions that remain compatible with vesicle handling (34). Likewise, immuno-affinity chromatography studies have shown that marker-directed EV capture in a chromatographic format is feasible under process-compatible conditions (35). In this context, the present workflow should not be viewed as a direct numerical benchmark against these systems, but rather as a complementary strategy that combines a recombinant affinity scaffold, EV-relevant release screening, and quantitative process evaluation within one development framework.

Orthogonal characterization further supports selective enrichment of a distinct EV fraction. The most direct enrichment readout was obtained by nano flow cytometry, where the CD81-positive fraction increased from 40% of total particles in conditioned medium to 95% and 91% in the eluates. In parallel, the eluates showed a smaller mean particle size and a narrower size distribution than the feed, and AF4 confirmed a corresponding shift in particle elution behavior. Together, these findings indicate that affinity capture did not simply concentrate the feed but enriched a compositionally different EV fraction.

The super-resolution analysis provided complementary information on TP composition but should be interpreted with more caution than the nano flow cytometry data. In conditioned medium, CD81 was already the dominant TP signal, and the cumulative CD81 ratio increased further in the eluates. At the same time, the strongest enrichment evidence in the present study comes from nano flow cytometry, whereas the super-resolution data are more informative for changes in TP composition and cluster organization than for absolute enrichment magnitude. It is therefore justified to state that the process enriched a CD81-positive and size-shifted EV fraction, but stronger conclusions regarding EV subclass identity or biological function would require additional evidence.

The affinity-enriched fractions also remained largely sensitive to Triton™ X-100 treatment, with particle counts decreasing by 83.6% in conditioned medium and by 90.6% and 87.9% in the two eluate samples, respectively. This supports the interpretation that the quantified particles were predominantly membrane-enclosed vesicular structures rather than detergent-resistant non-vesicular background. In addition, AF4 showed a distinct early protein peak in conditioned medium that was no longer present as a separate peak in the eluate, which is consistent with substantial depletion of non-associated soluble protein. At the same time, the remaining UV280 signal in the eluate co-eluted with the particle fraction and therefore cannot be interpreted as exclusively intended EV cargo without further analytical support.

The current arginine stability data are encouraging in that particle counts were maintained after exposure to the selected elution conditions. However, particle number alone is not sufficient to claim preserved biological activity.

Several limitations of the present study should be acknowledged clearly. Experimental validation is currently restricted to one target and one producer system, namely CD81-positive EVs from HEK293 cells. The Nanofitin® concept is in principle extensible to other surface markers, but transferability across targets remains a platform perspective rather than an experimentally demonstrated feature of the present work. The current column format was evaluated at 1 mL scale, which is appropriate for this stage of development but does not yet establish EV-specific scale-up behavior. In addition, the absence of a classical breakthrough profile prevented direct determination of a conventional DBC metric, so the measured dynamic binding should be interpreted as achieved binding under the tested conditions rather than maximal column capacity. Although NF06 showed favorable regeneration behavior in static BLI experiment, future development should include dynamic stability and reusability studies, ligand leaching assessment, and robustness testing across additional EV sources and feed compositions. In this context, prior reports on Nanofitin®-based affinity media in non-EV settings support the general suitability of the scaffold for chromatographic applications, but do not replace EV-specific validation (36, 22). Even with these limitations, the present data support the view that ligand-engineered affinity chromatography can provide a practical route toward marker-defined EV purification workflows that combine selectivity, controlled release, and process-compatible operation.

## Materials and Methods

### Selection of Anti-CD81 Nanofitin® Binders by Ribosome Display

A naïve Nanofitin® library based on the Sac7d scaffold was assembled by PCR using trinucleotide mutagenesis at the binding interface while excluding cysteine and proline to preserve scaffold integrity (37, 18). A human CD81 large extracellular loop (LEL; Phe113-Lys201) Fc chimera (R&D Systems, Minneapolis, MN, USA) was immobilized on Protein A/G magnetic beads (Cytiva, Marlborough, MA, USA) and used as target for ribosome display. Selection was performed for four consecutive rounds at 4 °C with progressively increased washing stringency as described in (38, 37, 18). Enrichment of CD81-reactive binders was monitored after each round by ELISA using CD81 LEL without Fc fusion as capture antigen. DNA recovered from round 4 was cloned into pQE30 to generate C-terminal GFP fusion constructs, transformed into *E. coli* DH5α LacIq cells (Invitrogen, Carlsbad, USA), and plated on 2YT plates containing 100 µg/mL ampicillin and 25 µg/mL kanamycin. Plates were incubated over night at 37 °C. Fluorescent positive clones were picked into 96-well plate containing 0.4 ml of 2YT medium with 100 μg/ml ampicillin, 25 μg/ml kanamycin and 1% glucose. After overnight shaking at 700 rpm, at 37°C preculture was transferred into fresh media and induced after 3 hours with β-D-1-thiogalactopyranoside (IPTG, 0.5 mM final concentration) further incubation at 700 rpm, 30°C overnight. Glycerol was added to the remaining pre-culture and stored at −80°C. Expression was stopped through centrifugation for 20 min at 2,000 x g and cell pellets were lysed with 1× BugBuster® reagent (MilliporeSigma, Burlington, MA, USA). After addition of 350 µL of TBS buffer (20 mM Tris-HCl, 150 mM NaCl, pH 7.4), lysates were clarified by centrifugation at 2,000 × g for 20 min. The resulting supernatants were used for fluorescence measurements and ELISA screening of individual clones.

### Expression and Purification of Candidate Nanofitins®

For pre-characterization of selected clones prior to biolayer interferometry (BLI), GFP-fused Nanofitin® candidates were expressed in 24-well format. Glycerol stocks from the initial 96-well screening were used to inoculate pre-cultures in 96 deep-well plates containing 800 µL of 2YT medium supplemented with 100 µg/mL ampicillin, 25 µg/mL kanamycin, and 1% glucose. After overnight incubation at 37 °C with shaking at 700 rpm, 200 µL of each pre-culture was transferred into duplicate 24-well plates containing 4 mL of 2YT medium supplemented with 100 µg/mL ampicillin, 25 µg/mL kanamycin, and 0.1% glucose per well. Cultures were incubated for 4 h at 37 °C and 700 rpm, induced with 50 µL of 40 mM IPTG (final concentration 0.5 mM), and expressed overnight at 30 °C with shaking at 700 rpm. Cells were harvested by centrifugation at 2,000 × g for 20 min at room temperature, and cell pellets were frozen at −80 °C for at least 1 h. Pellets were lysed by addition of 250 µL of 1× BugBuster® reagent (MilliporeSigma, Burlington, MA, USA) prepared in TBS buffer (20 mM Tris-HCl, 150 mM NaCl, pH 7.4) together with 0.125 µL DNase solution (10 mg/mL), followed by incubation for 1 h at room temperature with shaking. Lysates were clarified by centrifugation at 2,000 × g for 20 min at 4 °C. His-tagged proteins were captured on Ni-NTA 96-well plates (His-Pur™, Thermo Fisher Scientific, Waltham, USA) pre-equilibrated with TBS, washed sequentially with TBS, TBS containing 0.5% Triton™ X-100, and TBS, and eluted twice with 250 µL imidazole buffer (20 mM Tris, 500 mM NaCl, 500 mM imidazole, pH 7.4). Protein concentrations were determined by UV/Vis scan from 250 to 450 nm using a Lunatic spectrometer (Unchained Labs, Pleasanton, CA, USA).

For larger-scale production of C-terminal cysteine Nanofitin® variants, pET21 plasmids encoding the selected ligands were transformed into *E. coli* BL21-Gold cells (Agilent, Santa Clara, USA). Pre-cultures were prepared in 10 mL of 2YT medium supplemented with 100 µg/mL ampicillin and 1% glucose and incubated overnight at 37 °C with shaking at 600 rpm. Main cultures were grown in 500 mL flasks containing 200 mL of 2YT medium supplemented with 100 µg/mL ampicillin and 0.1% glucose until the optical density at 600 nm reached 0.8–1.0. Expression was induced by addition of 100 µL of 1 M IPTG and continued for 4 h at 30 °C. Cells were harvested by centrifugation at 3,220 × g for 30 min at room temperature, resuspended in 9 mL lysis buffer (20 mM Tris, 500 mM NaCl, 25 mM imidazole, pH 7.4), and stored overnight at −80 °C. After thawing, 1.5 mL of 10× BugBuster® reagent (MilliporeSigma, Burlington, USA), 7.5 µL DNase I (10 mg/mL), and 4.5 mL lysis buffer were added, followed by incubation for 15 min at room temperature with shaking. Clarified lysates were loaded onto Ni-NTA resin (300 µL settled resin; MilliporeSigma, Burlington, USA) in gravity-flow columns pre-equilibrated with ultrapure water and lysis buffer. The resin was washed with lysis buffer, lysis buffer containing 0.1% Triton™ X-100, and lysis buffer, and proteins were eluted with 1.5 mL elution buffer (20 mM Tris, 500 mM NaCl, 250 mM imidazole, pH 7.4). For each elution fraction, 167.5 µL of 100 mM EDTA was added. To improve recovery, the stored flow-through was reloaded once after re-equilibration of the resin with lysis buffer. Eluate concentrations were determined by UV/Vis scan from 250 to 450 nm using a Lunatic spectrometer (Unchained Labs, Pleasanton, CA, USA). Selected clones were sequenced, and alignment of mutated positions within the Nanofitin® binding site was used to identify related sequence families.

### Enzyme-linked Immunosorbent Assay (ELISA)

High-binding 96-well microplates (Nunc™, Thermo Fisher Scientific, Waltham, USA) were coated with 100 µL per well of CD81 LEL protein (without Fc fusion) diluted to 5 µg/mL in TBS buffer (20 mM Tris-HCl, 150 mM NaCl, pH 7.4). Plates were incubated for 1 h at RT 600 rpm, then washed three times with 300 µL/well of 1x TBS buffer. The wells were blocked with 300 µL/well 1x TBS containing 0.5% (w/v) bovine serum albumin (BSA) and incubated for 1 h at RT at 600 rpm, followed by four washes with 300 µL/well 1x TBS containing 0.1% Tween-20 (TBS-T). 100 µL/well of appropriately diluted Nanofitin® in TBS-T were incubated for 1h at RT while shaking at 600 rpm and subsequently washed four times with 300 µL of TBS-T. 100 µL RGS-HisTag antibody horseradish peroxidase conjugate (Qiagen, Hilden, Germany), diluted 1:4000 in TBS-T, was added per well and incubated for 1 h at RT while shaking at 600 rpm. After four consecutive washing steps with 300µL of TBS-T, signal was developed by the addition of 100µL of 1 mg/mL freshly prepared tetramethylbenzidine (TMB) substrate (100µL of 10 mg/mL of TMB diluted in 0.1 M citrate/acetate, pH 6.0, containing 4µL 30% H_2_O_2_) after brief mixing at 600 rpm and 2-5 min of incubation, reactions were stopped with 100 µL/well of 1 M HCl, plates were sealed with optical film, and absorbance at 450 nm was measured using a microplate reader (Varioskan, Thermo Fisher Scientific, Waltham, USA).

### Biolayer Interferometry (BLI)

Binding kinetics and elution-related dissociation were analyzed on an Octet R8 instrument (Sartorius, Göttingen, Germany) at 25 °C and 1000 rpm. GFP-fused candidates were captured on streptavidin biosensors via a biotinylated anti-GFP Nanofitin®, whereas C-terminal cysteine variants were biotinylated using EZ-Link Maleimide-PEG2-Biotin (Thermo Fisher Scientific, Waltham, USA) and loaded directly onto streptavidin biosensors. After baseline acquisition in standard assay buffer (TBS with 0.1% BSA and 0.02% Tween-20), sensors were exposed to CD81 LEL at the indicated concentrations for 180 s, followed by 180 s dissociation. For elution screening, the dissociation step was performed in the indicated elution buffers and corrected using identically loaded reference sensors without target. For ligand robustness testing, loaded sensors were cycled 20 times through target binding, regeneration, and 10 min exposure to either 0.1 M NaOH or 0.1 M glycine, pH 2.0. Data were analyzed using Octet Data Analysis HT software V13.0 using the 1:1 model after reference subtraction.

### Immobilization of Nanofitins® on Eshmuno® Epoxy Resin

Nanofitins® were immobilized on Eshmuno® epoxy-activated resin (Merck KGaA, Darmstadt, Germany) by mixing the ligand stock solution with the resin at a ratio of 20:1 (mL/g) in phosphate buffer, pH 9.0, after prior reduction of the Nanofitins® with 10 mM Dithiothreitol (DTT). The coupling reaction was carried out for 6 h at 40 °C under agitation. The resin was then vacuum-filtered and washed with coupling buffer. Remaining free epoxy groups were deactivated by addition of amines and adjusted to a final concentration of 10% (v/v). The suspension was incubated overnight at room temperature under agitation. After incubation, the resin was vacuum-filtered and washed with deactivation solution. To remove non-covalently bound ligand, the resin was subjected to sequential washing with 0.1 M NaOH and 0.1 M glycine, pH 3.0, followed by extensive washing with storage solution. The final resin was stored as a 50% (v/v) slurry in storage solution (150 mM NaCl, 20% ethanol).

### Column Packing

For column packing, resin slurry was adjusted to 50% (v/v) in wash buffer and settled using vacuum, followed by packing with a 10% compression rate. Columns were initially cleaned with 10 CV of 0.1 M NaOH, followed by 10 CV of 50 mM glycine, pH 3.0, and 10 CV of wash buffer at a flow rate of 1 CV/min. After initial cleaning, columns were stored in storage solution.

### Static Binding and Elution Screening with Recombinant CD81 and EVs

Static binding assays were performed in MultiScreen HTS-DV filter plates (0.65 µm pore size, MilliporeSigma) using Nanofitin®-coupled resin equilibrated in wash buffer (50 mM Tris-HCl, 125 mM NaCl, pH 7.4). For recombinant target screening, 10 µL settled resin was incubated with 200 µL recombinant CD81 LEL (0.5 mg/mL) for 120 min at room temperature with shaking. Flow-through and wash fractions were collected by centrifugation, and bound protein was eluted in two sequential steps using the indicated elution buffers. Protein concentrations in all fractions were determined from A280 using a calibration curve generated with recombinant CD81 standards. For EV screening, the settled resin volume was increased to 30 µL per well. AEX-purified HEK293-derived EVs were diluted to 2 × 10^10^ particles/mL in wash buffer, and 200 µL was applied per well. Binding, washing, and elution were performed as described for recombinant CD81. CD81-positive EVs in flow-through, wash, and elution fractions were quantified by CD81 ELISA as described below.

### HEK293 Cell Culture, Conditioned Medium Harvest, and TFF Concentration

Suspension-adapted HEK293 cells (cat. VP002, MilliporeSigma) were cultured in chemically defined, serum-free Cellvento 4HEK medium supplemented with 6 mM L-glutamine in shaking flasks at 37 °C and 5% CO_2_. Cultures were seeded at 5 × 10^5^ viable cells/mL and harvested after 48 h at 5 × 10^6^ viable cells/mL and 99% viability. Cells and large debris were removed by sequential centrifugation at 500 × g for 5 min and 2000 × g for 20 min, both at 4 °C. The resulting conditioned medium was aliquoted and stored at −80 °C until further processing. For preconcentration, thawed conditioned medium was clarified through a 0.22 µm SteriCup® vacuum filter (MilliporeSigma) and concentrated 10-fold by tangential flow filtration using Pellicon® XL 50 cassettes with a 100 kDa molecular weight cutoff (MilliporeSigma). TFF was performed at room temperature using a feed flow rate of 35 mL/min while maintaining the transmembrane pressure between 1.5 and 2.0 bar.

### Affinity Chromatography

TFF-concentrated conditioned medium was processed on an ÄKTA pure 25 M system (Cytiva) equipped with a fraction collector and a 1 mL Eshmuno® anti-CD81 Nanofitin® column. Columns were equilibrated with 10 column volumes (CV) of wash buffer (50 mM Tris, 125 mM NaCl, pH 7.4). Feed was loaded at 0.33 CV/min, corresponding to a residence time of 3 min, followed by a 10 CV wash at 1 CV/min. Bound material was eluted with 10 CV of elution buffer (1 M arginine, 50 mM Tris, pH 10). Columns were cleaned in place with 10 CV 0.1 M NaOH, re-equilibrated with wash buffer, and stored in 150 mM NaCl containing 20% ethanol. Flow-through, wash, and elution fractions were collected for subsequent analysis. EV-containing elution fractions were pooled, buffer-exchanged three times against wash buffer using Amicon ultrafiltration devices (3 kDa cutoff, 4500 × g, 20 min per cycle), and concentrated for downstream analysis. UV absorbance at 260 and 280 nm and inline multi-angle light scattering were recorded throughout each run.

### EV Characterization

#### Nano Flow Cytometry (nFCM)

Single-particle measurements were performed on a NanoAnalyzer instrument (NanoFCM Inc., Xiamen, China) equipped with 488 and 640 nm lasers and avalanche photodiode detectors for side scatter and fluorescence detection. Particle-free Milli-Q water was used as sheath fluid, and measurements were acquired for 60 s at an operating pressure of 1 kPa. Samples were diluted in 25 mM HEPES, pH 7.4, to yield 1500-12,000 detected events per acquisition. Milli-Q water and HEPES blanks were required to remain below 100 particles min^−1^. Absolute particle counts were calibrated using 250 nm silica standards (NanoFCM Inc., Xiamen, China), and particle size was estimated from a four-modal silica calibration mixture (NanoFCM Inc., Xiamen, China) according to the manufacturer’s workflow. For CD81 phenotyping, samples were first adjusted to 5 × 10^9^ to 10^10^ particles/mL in particle-free PBS and incubated for 30 min in the dark with FITC-conjugated anti-CD81 antibody (clone 5A6, BioLegend, San Diego, USA) at a final concentration of 10 mg/mL. Matched FITC-IgG controls and antibody-only controls were processed in parallel. After staining, samples were diluted in particle-free HEPES buffer to the measurement range and acquired in duplicate. Gates for CD81-positive events were defined in nFCM Professional Suite v3.0. To assess detergent sensitivity, samples were mixed 1:1 with 10% Triton™ X-100 (final concentration 5%), incubated for 30 min, and measured after dilution to the same final factor as untreated controls.

#### Enzyme-linked immunosorbent assay (ELISA) for CD81+ EV quantification

For the quantification of intact EVs, a sandwich ELISA against human CD81 (CD81-Capture Human Exosome ELISA Kit, FUJIFILM Wako Pure Chemical Corporation, Osaka, Japan) was implemented. The ELISA was performed according to the manufacturer’s instructions. In brief, prewashed multiwell plate, containing immobilized antibody, was incubated for 2 h with samples and subsequently washed. Biotinylated anti-CD81 was incubated for 1h and exchanged for streptavidin conjugated with horseradish peroxidase, after washing. After last wash step, Tetramethylbenzidine (TMB) was added as substrate and the reaction was stopped after 30 minutes of incubation. Plates were measured on a microplate reader (Infinite M Nano+, TECAN, Männedorf, Switzerland) at 450nm and 620nm (correction). TFF concentrated CM, used also as load for affinity chromatography, was used for absolute quantification through the generation of a standard curve. CM was serially diluted, and samples were diluted accordingly to fit into the range of the standard curve. Absolute CD81+ EV value for the Load was determined through nFCM staining. Graphpad Prism software V10.2.1 was used for the interpolation of the unknown EV concentration using a third order polynomial regression curve and log-log plot. All measurements were done in duplicates.

#### Asymmetric-flow field-flow fractionation (AF4)

Fractionation and detection of particles was performed using an Eclipse® AF4 system (Wyatt Technology Santa Barbara, USA) coupled with an Agilent HPLC 1260 system (Agilent Technologies Inc, Santa Clara, USA) equipped with autosampler, pump, UV-, refractive index-, MALS- and Quasi-Elastic Light Scattering-detector. For separation, a short channel (144 nm length, 400 µm thickness) with regenerated cellulose membrane,10kDa molecular weight cut-off was used. Running buffer comprised of 20 mM Na2HPO4, 350 mM NaCl and 250 ppm sodium azide. All samples were injected twice. The method consisted of 2 minutes equilibration at 2 mL/min, injection start with 1 minute focusing at 2 mL/min, followed by 0.2 mL/min for 7 minutes and afterwards again focusing at 2 mL/min for 7 minutes. Elution started with constant elution for 5 minutes at 2 mL/min, followed by an exponential gradient at 0.05 mL/min (20 min-slope 6) and constant elution again for 20 minutes at 0.05 mL/min. Channel flow was set to 1 mL/min and detector flow at 0.5 mL/min. Data was acquired and evaluated using Astra Software V8.1.1.12 using online particle and number density templates. Running buffer was measured as blank for baseline subtraction.

#### Super Resolution Microscopy

For glas functionalization, 1.5H cover glasses were first cleaned in a methanol/HCl (1:1) bath. Then, they were rinsed with methanol, dried and exposed to ozone. Silanization was carried out with 1 mg/ml silane-PEG-NHS (5kDa PEG, Nanocs Inc., New York, USA) in 95% (v/v) for 20 minutes at RT. Unreacted silanes were washed out with two consecutive washes of 95% (v/v) ethanol and Milli-Q water. For EV capture, sample pH was adjusted by adding 1/10 volume of 1M sodium bicarbonate. Subsequently, EVs were incubated on the functionalized surface over night at 4°C. Unbound EVs were then washed out with Tris-buffered saline containing 0.02% Tween-20 (TBS-T), followed by a 4% formaldehyde fixation for 10 min at RT. After two TBS-T washes, the sample was blocked with 10% (w/v) BSA for 45 minutes. After blocking, epitopes were detected with an antibody cocktail (anti-CD9, 488 nm; anti-CD63, 561 nm; anti-CD81, 647 nm; ONI, Oxford, UK; 3% BSA) for 50 min at RT. Unbound antibodies were washed out with three subsequent TBS-T washes and the remaining antibodies were fixed in place with 4% formaldehyde for 10 min. Following two TBS-T washes, the sample was pre-incubated for 10 min at RT with dSTORM sulfite buffer (30 mM DTT, 30 mM NaSO3, 65 mM triethylenediamine, in 1x PBS). After pre-incubation the imaging buffer was refreshed and imaging was performed.

Imaging was conducted in a NanoImager (ONI, Oxford, UK), which was set to 32°C under TIRF illumination. Fluorophore blinking was induced with high laser power, and 1000 frames were imaged per channel with a 30 ms exposure time. Fluorophore localization, image reconstruction and cluster analysis were subsequently carried out with the CODI analysis platform (ONI, Oxford, UK). A cluster was defined as an EV if >10 localizations (irrespective of marker) were observable within a 70 nm search radius and an EV was considered positive for a respective marker if >5 localizations of the specific marker were observable.

### Impurity Assays

#### Host cell protein (HCP) ELISA quantification

HEK293 host cell protein (HCP) concentrations in chromatographic fractions were determined using a HEK293 HCP ELISA kit (F560s-ELISA Kit, Gygnus Technologies, Leland, USA) according to the manufacturer’s instructions. Briefly, appropriately diluted samples were dispensed into the antibody-containing microplate and incubated for 90 min. Plates were washed, developed with TMB substrate for 30 min, and the reaction was stopped according to the kit protocol. Absorbance was measured at 450 nm with 650 nm correction using an Infinite M Nano+ microplate reader (TECAN, Männedorf, Switzerland). HCP concentrations were interpolated in GraphPad Prism V10.2.1 using a third-order polynomial regression on a log-log plot. All measurements were performed in duplicate.

#### Total protein quantification by bicinchoninic acid (BCA) assay

Total protein concentrations were determined using the Pierce™ BCA Protein Assay Kit (Thermo Fisher Scientific, Waltham, MA, USA) according to the manufacturer’s instructions. Briefly, measurements were performed using the microplate procedure with a sample-to-working reagent ratio of 1:8. Samples and BSA standards (25-2000 µg/mL) were diluted in equilibration/wash buffer to appropriate concentrations and incubated with working reagent for 30 min at 37 °C. Absorbance was measured at 562 nm using an Infinite M Nano+ microplate reader (TECAN, Männedorf, Switzerland). Protein concentrations were determined in GraphPad Prism V10.2.1 using a fourth-order polynomial fit. All measurements were performed in duplicate.

#### Double-stranded DNA (dsDNA) quantification by PicoGreen™ assay

dsDNA concentrations in chromatographic fractions were determined using the Quant-iT™ PicoGreen™ dsDNA Reagent and Kit (Invitrogen, Carlsbad, CA, USA) according to the manufacturer’s instructions. Low-range standards and samples were diluted in TE buffer, mixed with freshly prepared working solution in a microplate, and incubated for 5 min in the dark. Fluorescence was measured using an Infinite M Nano+ microplate reader (TECAN, Männedorf, Switzerland) at 480 nm excitation and 520 nm emission. Sample dsDNA concentrations were determined using a linear standard curve generated in Microsoft Excel. All measurements were performed in duplicate.

### Arginine Stability Assay

Purified EVs were buffer exchanged by ultrafiltration (Amicon® 3 kDa cut-off, 3 × 4,500 × g, 20 min) into 0.1, 0.5, or 1 M arginine containing 50 mM Tris, pH 10, or into PBS as control, and incubated for 0, 0.5, 1, 4, or 24 h at 4 °C. After each incubation time, EVs were buffer-exchanged three times into equilibration/wash buffer using the same ultrafiltration protocol, and particle concentration was measured by nFCM. For each time point, data obtained for the different arginine concentrations were normalized to the corresponding PBS control.

### Statistical Analysis and Calculations

Results are presented as mean ± standard error of the mean (SEM), unless stated otherwise. Statistical analysis was performed using GraphPad Prism 10 V10.2.1.

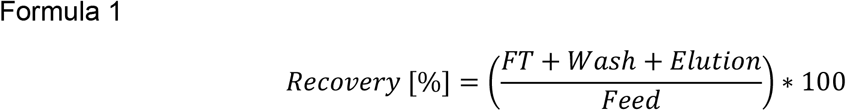

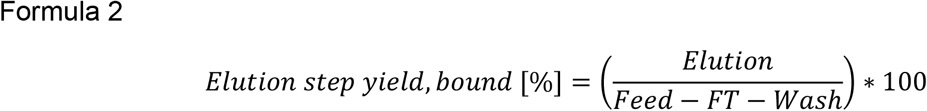

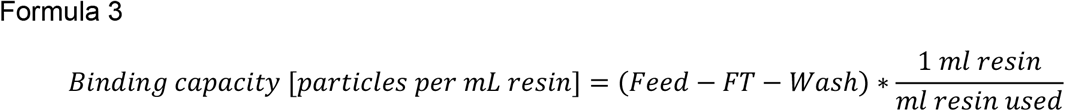

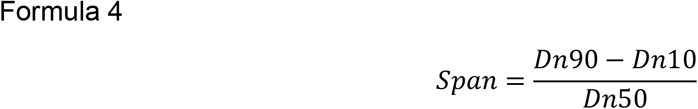

Dn represents the particle diameter corresponding to 10% (D10), 50% (D50), or 90% (D90) of the cumulative particle number distribution.

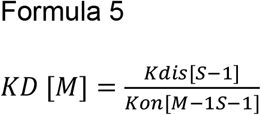

## Author Contributions

CG conceived, designed and performed experiments regarding Nanofitin® screening, characterization, lead candidate selection and analyzed the data. KB established the super-resolution microscopy experiments and FC performed and analyzed the data. TB performed AF4 analysis. LFK designed and performed the remaining experiments and analyzed the data. EW analyzed the data and contributed to study conception. MJS conceived the study and supervised the project. LFK and MJS drafted the manuscript including participation from CG. CG and EW contributed to manuscript revision and editing. All authors have read and agreed to the final version of the manuscript.

## Acknowledgement

We want to thank Dr. Romas Skudas and Dr. Mathieu Cinier for their scientific advisory regarding Nanofitin® design and selection. We want to thank Dr. Peter Menstell for his support in the SBC expermients and thank Daniel Schneider for performing the resin coupling.

Figure 1 was created in BioRender. Saul, M. (2026) https://BioRender.com/noi7csy. Figure 3 was created in Biorender. Saul, M. (2026) https://BioRender.com/b5cppyw

