## Supplementary Figures for "Nanofitin®-Engineered Affinity Chromatography for Marker-Defined Extracellular Vesicle Enrichment in Scalable Downstream Processing"

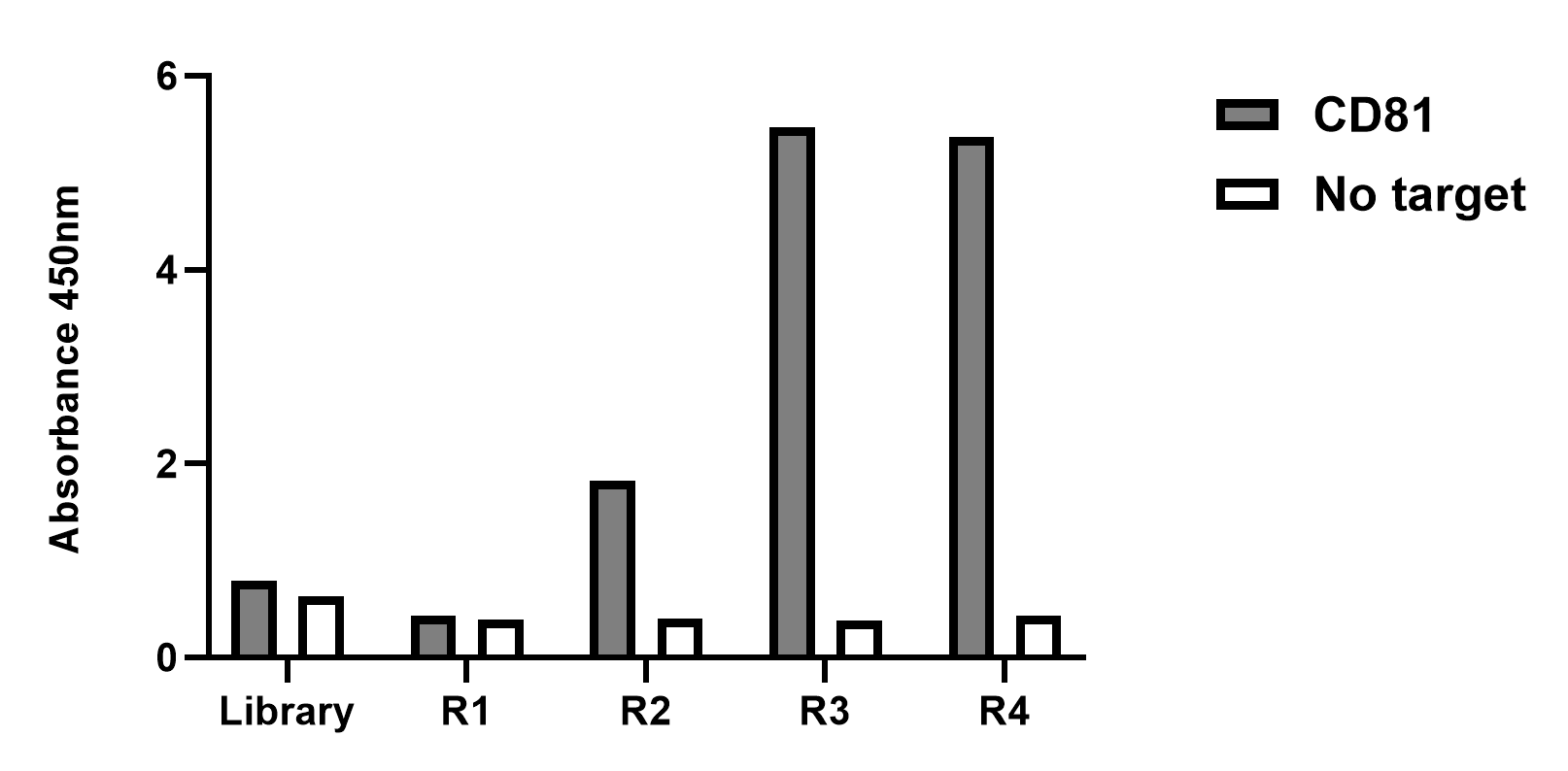

Supplementary Figure 1. Enrichment of CD81-binding Nanofitin® candidates through ribosome display selection. Screening of CD81-binding capabilities of naïve library and pooled candidates after each selection round against unspecific binding, quantified using enzyme-linked immunosorbent assay. Data represents single measurement.

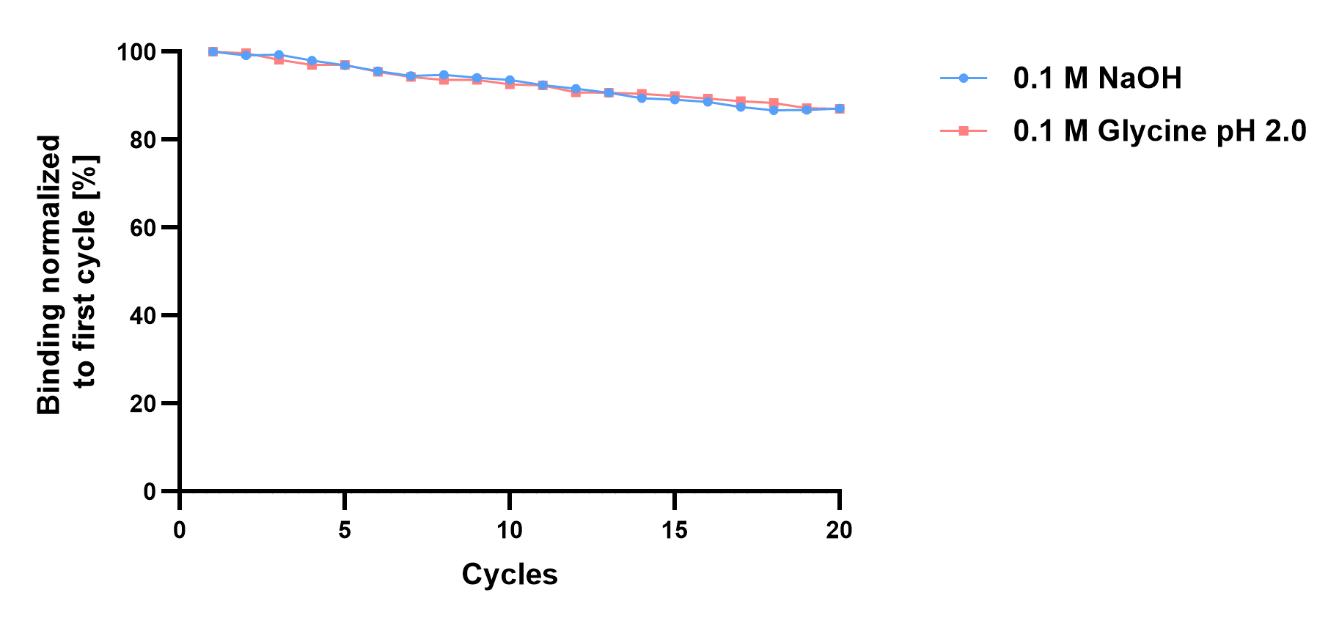

Supplementary Figure 2. Stability of Nanofitin® candidate NF06 towards caustic and low pH conditions. Retention of binding capabilities of NF06 over 20 cycles in 0.1 M NaOH and 0.1 M Glycine (pH 2), data normalized to control sensor (Tris buffered saline solution, pH 7.2) and first cycle. Data represents single measurement.

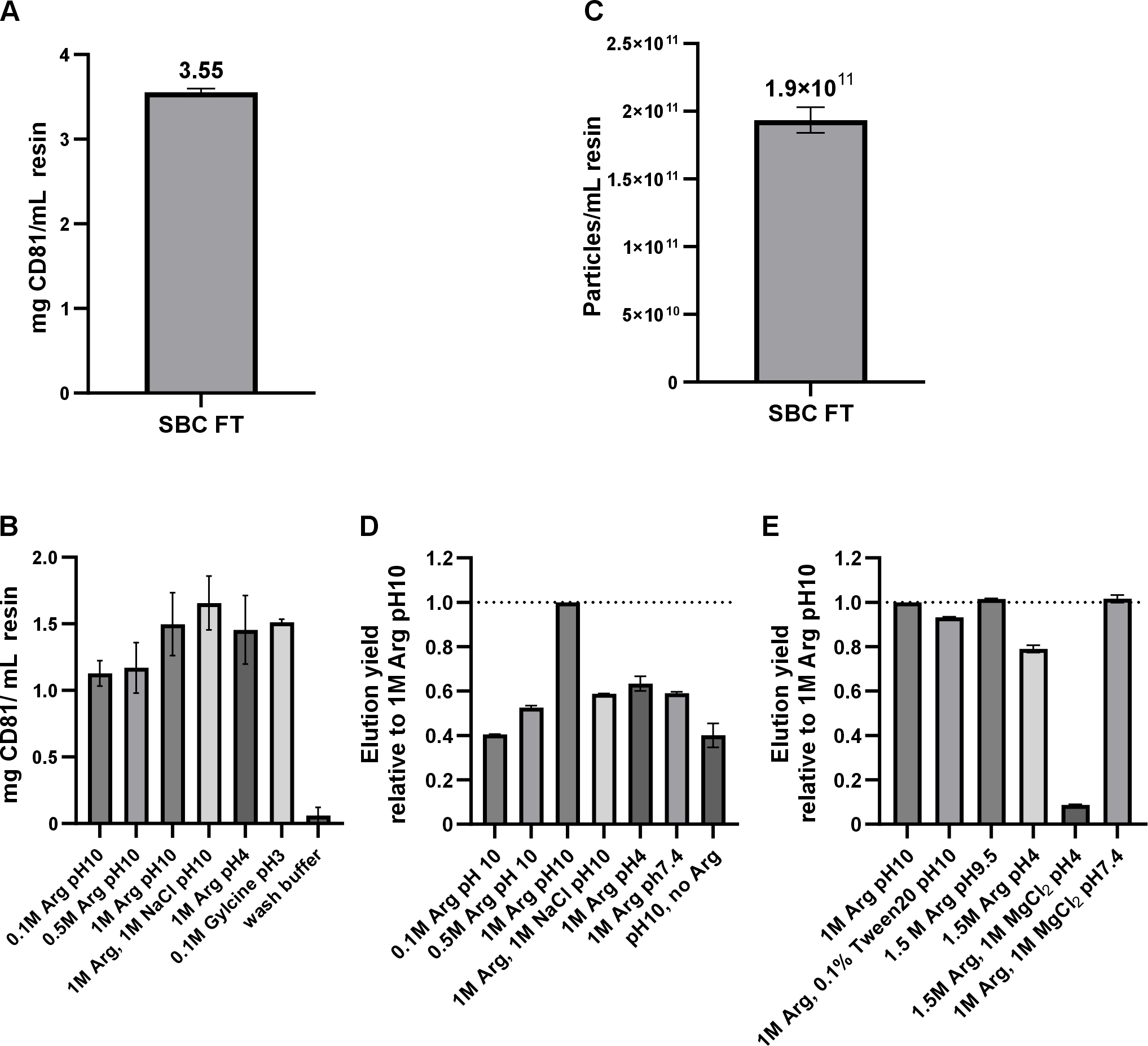

Supplementary Figure 3. Static binding capacity and elution performance of Nanofitin®-coupled resin with recombinant CD81 protein and CD81-positive extracellular vesicles. (A) Static binding capacity (SBC) based on Flow-through of NF06-coupled resin with recombinant large extracellular (LEL) loop of CD81. (B) Elution buffer screening of recombinant CD81 LEL bound to resin-coupled NF06 with elution buffers varying in composition. (C) Flow-through based SBC of NF06 with HEK293-derived extracellular vesicles (EV). (D) First elution buffer screening using HEK293-derived EVs to confirm previous protein-based elution, normalized on the elution of buffer condition containing 1M arginine at ph10. (E) Extended elution buffer composition screening using EVs, normalized on the elution of buffer condition containing 1M arginine at pH 10. Data represents mean +/- Standard Error of the Mean (SEM) of n=2.

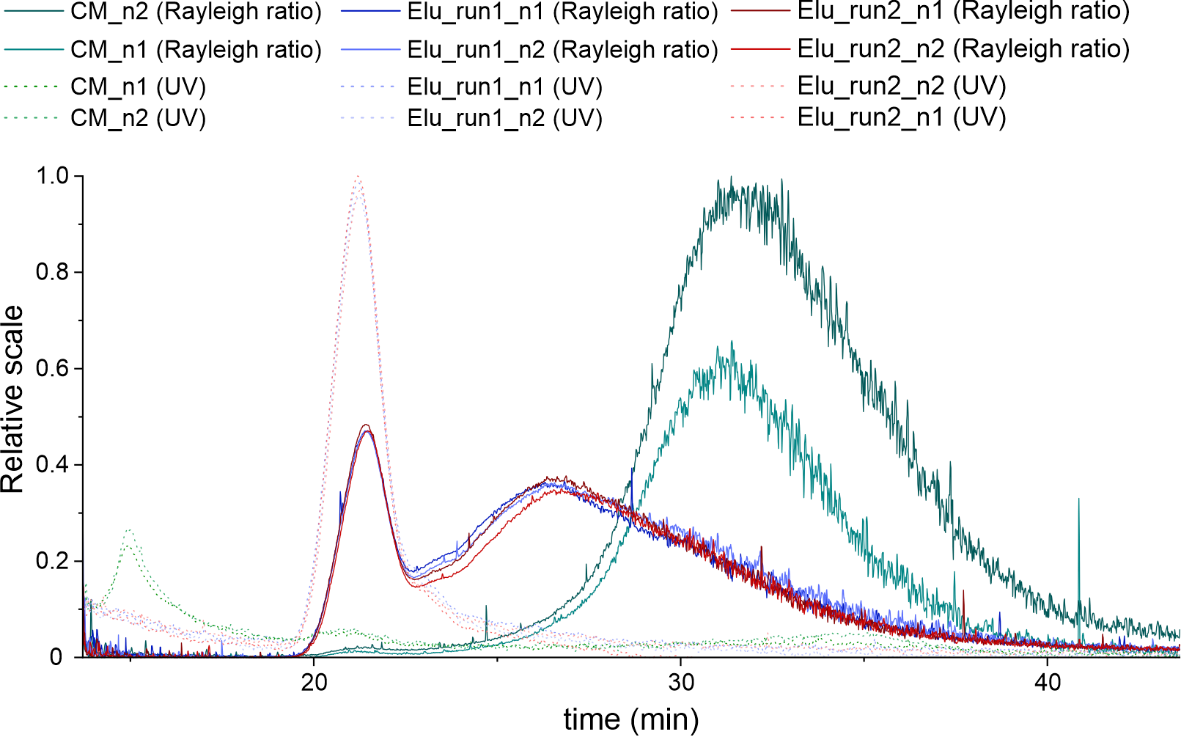

Supplementary Figure 4. Protein and particle profile of extracellular vesicles before and after affinity isolation. Asymmetric-flow field-flow fractionation chromatogram of conditioned media (CM) and Elution (Elu) of two NF06 affinity chromatographic runs with UV280 nm detection (dotted line) and multiangle light scattering (MALS) detection (continuous line), measured in two technical replicates. Data represented in relative scale to highest value.

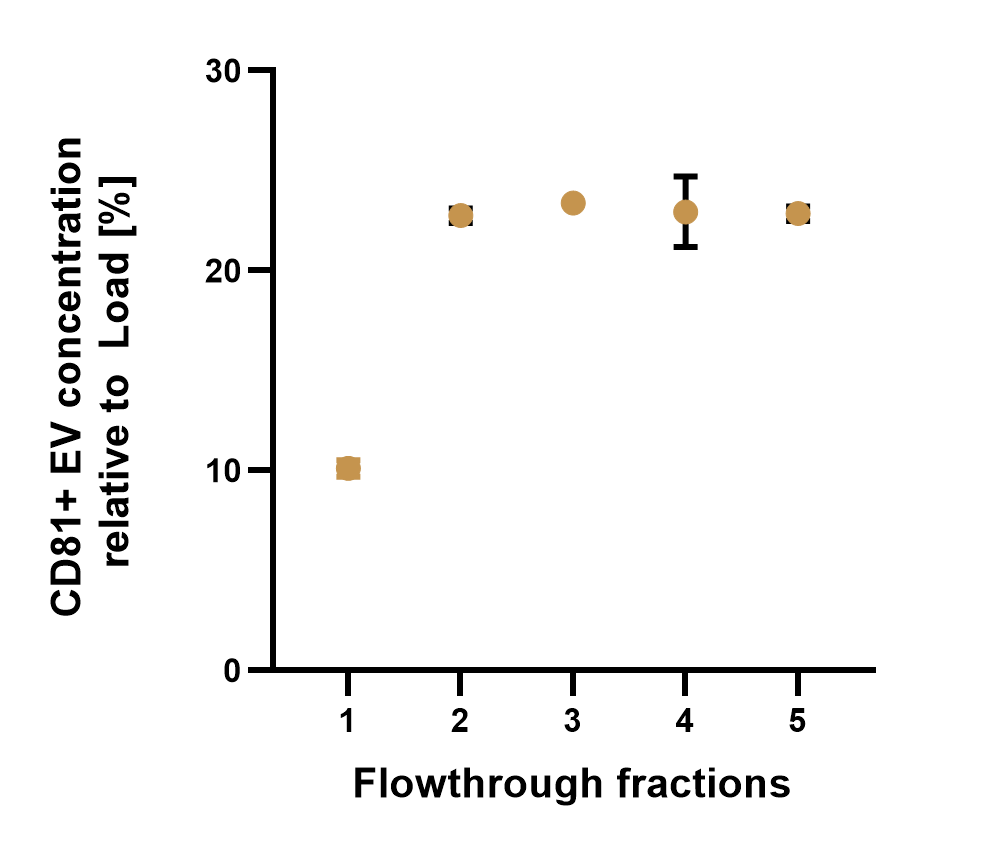

Supplementary Figure 5. Presence of CD81-positive extracellular vesicles (EV) in the Flow-through (FT) fractions during Nanofitin® affinity chromatography. Quantification of CD81-positive EVs through enzyme-linked immunosorbent assay (ELISA) in subsequent FT fractions from affinity chromatography using Eshmuno®-coupled Nanofitin® NF06 prototype column. Concentration was normalized to CD81-positive EV concentration of the Load. Data represents mean +/- Standard Error of the Mean (SEM) of n=2.

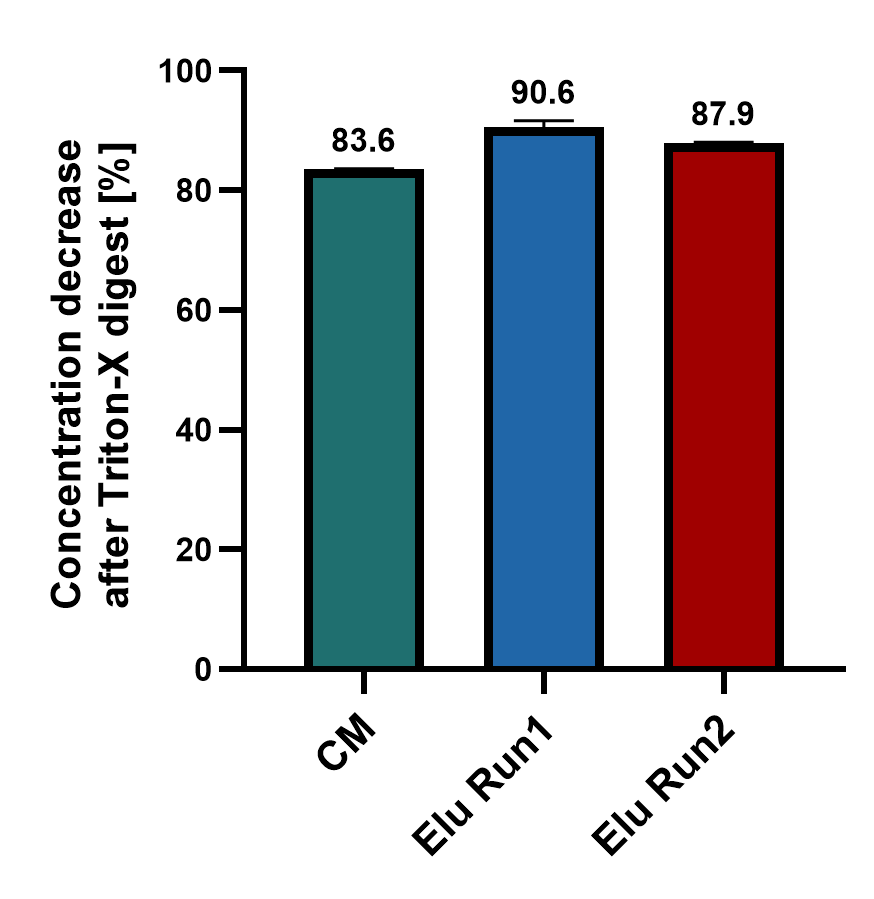

Supplementary Figure 6. Disruption of membrane enclosed extracellular vesicles (EV) through Triton X-100 digest. Ablation of membranous particles present in conditioned media (CM) and Elution (Elu) from NF06 affinity chromatography run 1 and 2 was quantified using nano flowcytometry after sample incubation with 5% Triton-X100 for 30 minutes. Reduction in particle concentration was calculated from untreated sample Data represents mean +/- Standard Error of the Mean (SEM) of n=2.

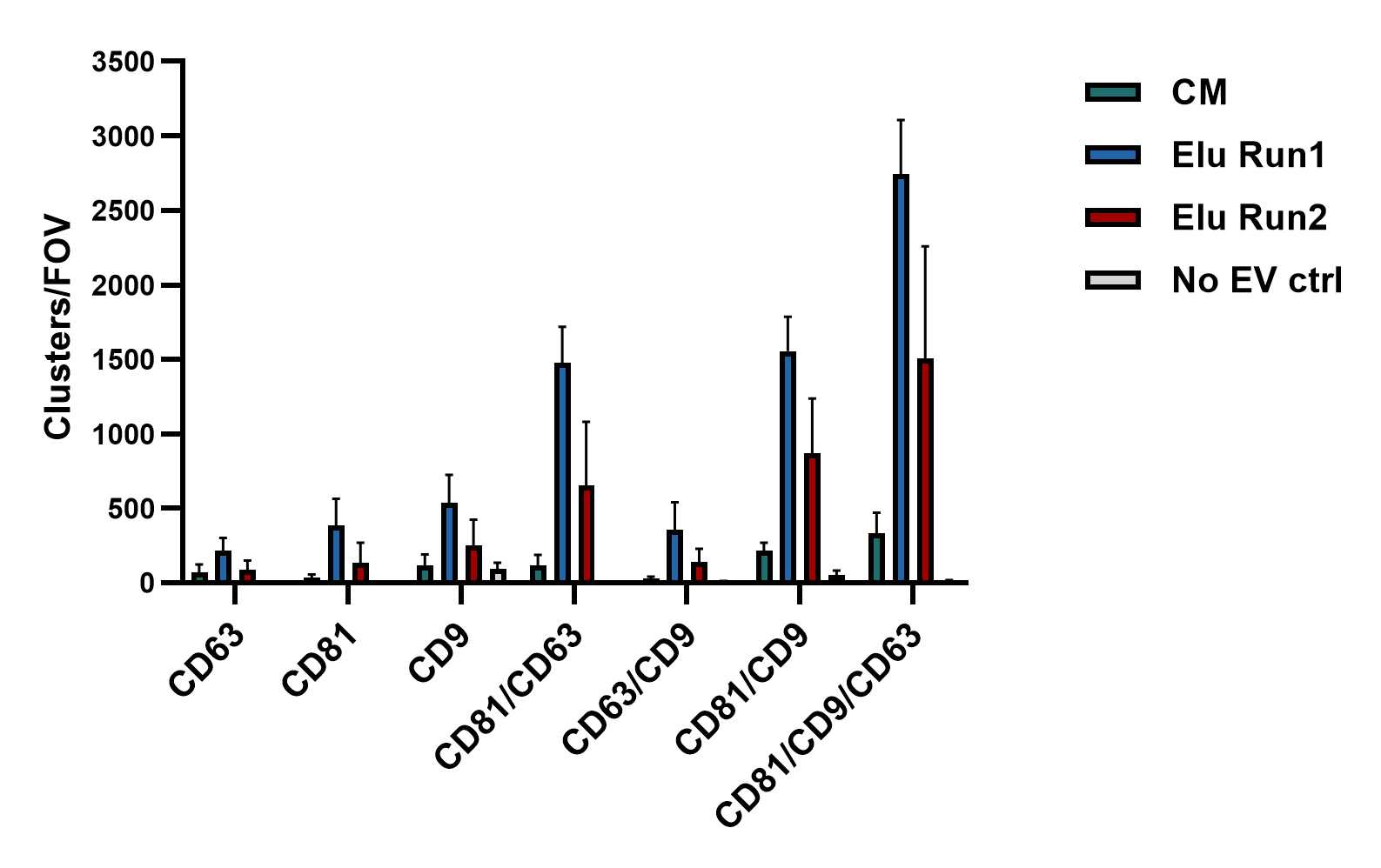

Supplementary Figure 7. Tetraspanin colocalization count of extracellular vesicles (EV). Super-resolution microscopy quantification of fluorescent clusters and their colocalization from the tetrapsanin (CD9, CD63, CD81) staining of EVs from HEK293 conditioned media (CM) and Elution (Elu) from two NF06 affinity chromatographic runs, as well as no EV control. Data represents mean +/- Standard Error of the Mean (SEM) of 5 technical replicates.

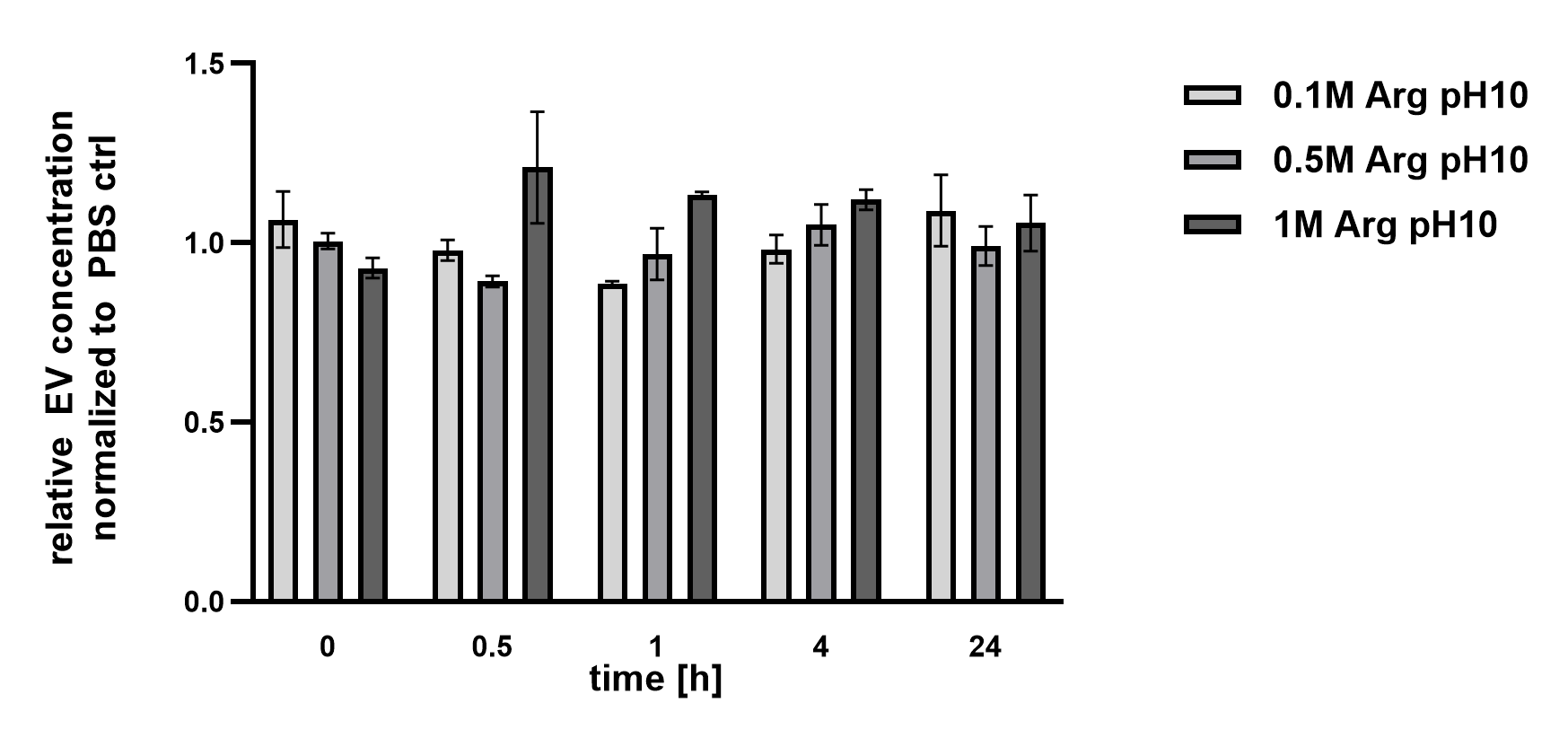

Supplementary Figure 8. Stability assessment of extracellular vesicles (EV) in arginine buffer. EV stability was assessed by quantifying EVs, using nano flowcytometry, after incubation in buffers containing 0.1-1M arginine at pH 10 for up to 24 hours at 4°C. Data was normalized on PBS control at same timepoint. Data represents mean +/- Standard Error of the Mean (SEM) of n=2.

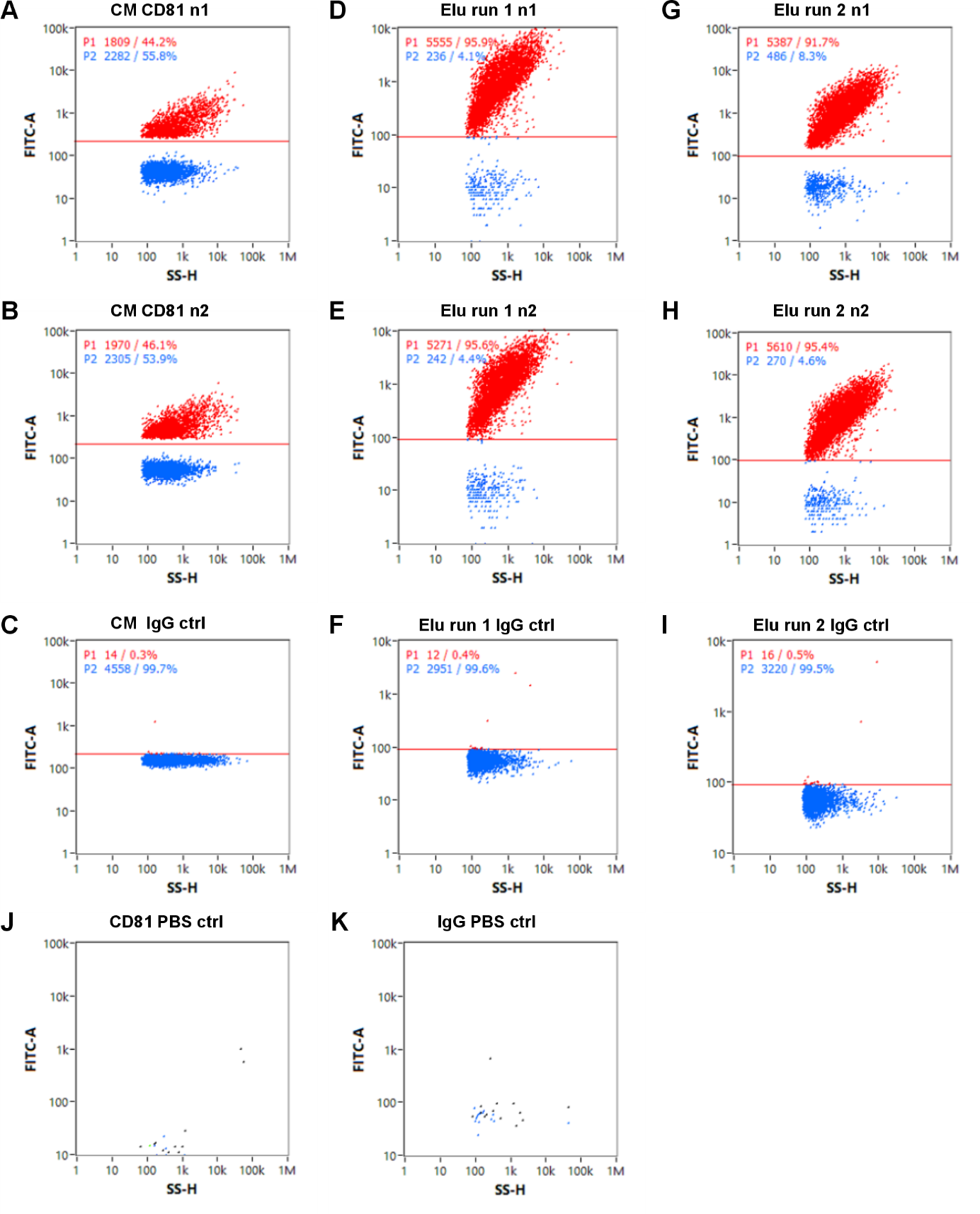

Supplementary Figure 9. Nanoflow cytometry stainings and controls of Conditioned Media (CM) and Elution (Elu) of the two NF06 affinity chromatographic runs. (A-C) CM measurements of FITC-labeled CD81 antibody staining with two technical replicates and gating strategy based on FITC-labeled IgG antibody. (D-F) Elu of run 1 measurement of FITC-labeled CD81 antibody staining with two technical replicates and gating strategy based on FITC-labeled IgG antibody. (G-I) Elu of run 2 measurements of FITC-labeled CD81 antibody staining with two technical replicates and gating strategy based on FITC-labeled IgG antibody. (J+K) PBS control of both FITC-labeled antibodies used for immunofluorescence staining.
